# Presynaptic development is controlled by the core active zone proteins CAST/ELKS

**DOI:** 10.1101/823542

**Authors:** Tamara Radulovic, Wei Dong, R. Oliver Goral, Connon I. Thomas, Priyadharishini Veeraraghavan, Monica Suarez Montesinos, Debbie Guerrero-Given, Kevin Goff, Matthias Lübbert, Naomi Kamasawa, Toshihisa Ohtsuka, Samuel M. Young

## Abstract

Many presynaptic active zone proteins have multiple regulatory roles that vary during distinct stages of neuronal circuit development. However, our understanding how presynaptic active zone proteins regulate synapse development during neuronal circuit maturation is in its early stages. Although CAST/ELKS are presynaptic active zone core proteins, their roles in synapse development in the mammalian central nervous system remain enigmatic. To unravel CAST/ELKS roles in glutamatergic synapse development, we analyzed how their loss during the early stages of circuit maturation impacted the calyx of Held presynaptic terminal development and function. We found a reduction in presynaptic surface area and volume, but an increase in active zone size. Additionally, we found a reduction in all presynaptic Cav2 channel subtype currents. Surprisingly, these changes did not impair synaptic transmission. We propose that CAST/ELKS are involved in pathways regulating presynaptic morphological properties and Cav2 channel subtype levels during early stages of neuronal circuit maturation.

## Introduction

The presynaptic active zone (AZ) is composed of a network of proteins with multiple roles that regulate synaptic function and the diversity of information encoding by neuronal circuits (Nusser, 2018). Many of these proteins are expressed as multiple isoforms with different regulatory domains that vary between different neuronal populations and stages of synapse development during neuronal circuit maturation (Ackermann et al., 2015). It is emerging that many neurological disorders are caused by alterations in neurodevelopment (Willsey et al., 2018) and mutations in presynaptic proteins can lead to neurological disorders (Cortes-Saladelafont et al., 2018). Therefore, it is important to understand how presynaptic AZ proteins regulate synapse maturation during neuronal circuit development.

The mammalian CAST/ELKS protein family are highly conserved core AZ proteins (Ohtsuka et al., 2002; Wang et al., 2002; Hagiwara et al., 2005) that are orthologs of *D. melanogaster bruchpilot* (Wagh et al., 2006) and *C. elegans elks* (Deken et al., 2005). They directly interact with Munc13 (Wang et al., 2009), Liprin-α (Ko *et al.*, 2003), Bassoon/Piccolo (Takao-Rikitsu *et al.*, 2004), RIMs (Ohtsuka et al., 2002; Wang et al., 2002), SAD-B kinase (Mochida et al., 2016) and the Cavβ4 subunit (Kiyonaka et al., 2012). Based on these numerous interactions, CAST/ELKS are implicated in regulating synapse development and function. Genetic studies on *bruchpilot*, at *Drosophila* T-bar synapses *in vivo*, demonstrated multiple roles regulating presynaptic active zone morphology, recruitment of voltage-gated calcium channels (VGCCs), and synaptic transmission (Kittel et al., 2006; Wagh et al., 2006; Fouquet et al., 2009). However, initial loss of function studies on *elks* at *C. elegans* en passant synapses found no changes in synaptic morphology (Deken et al., 2005). Genetic studies at mammalian retinal ribbon synapses *in vivo* which deleted both CAST/ELKS identified roles for CAST/ELKS regulation of synapse formation and AZ morphology (Hagiwara et al., 2018). However, deletion of both CAST/ELKS at the mature calyx of Held *in vivo*, a large Cav2.1 exclusive glutamatergic presynapse with many conventional AZs in parallel (Baydyuk et al., 2016) lead to no changes in presynaptic morphology or AZ ultrastructure (Dong et al., 2018). In addition, studies on *in vitro* cultured hippocampal deleted for both CAST/ELKS found no changes in AZ ultrastructure in GABAergic neurons (Liu et al., 2014) and in glutamatergic neurons (Held et al., 2016).

However, these previous studies at mammalian conventional AZ synapses did not analyze CAST/ELKS function in a developing native neuronal circuit *in vivo*. Therefore, it is unclear if CAST/ELKS roles in regulating synapse development and AZ ultrastructure is unique to ribbon synapses, or these roles are confined to a specific developmental stage of synapse maturation. In addition, although a role for CAST/ELKS was found for regulating Cav2.1 current levels and channel numbers (Dong et al., 2018), it is not known if CAST/ELKS regulates all presynaptic Cav2 subtype levels or if it solely Cav2.1 specific.

To address these questions, we investigated the role of CAST/ELKS in regulating synaptic function in the immature calyx of Held (P9-11), in the developing auditory brainstem. The calyx of Held is an excellent model system to study synapse development and function in a native neuronal circuit (Baydyuk et al., 2016), and presynaptic underpinnings of neuropsychiatric disorders (Di Guilmi et al., 2014; Montesinos et al., 2015; Wang et al., 2015; Zhang et al., 2017). During early development synaptic transmission at the calyx of Held relies on three subtypes of Cav2 channels which makes it ideal for studying effects of CAST/ELKS on specific Ca^2+^ subtypes. We used *Cast/Erc2* KO (*Cast^−/−^*), *Elk1s/Erc1* conditional (CKO) transgenic mice (Dong et al., 2018; Hagiwara et al., 2018) in conjunction with our ability to conditionally ablate ELKS expression in the calyx of Held (Dong et al., 2018) and demonstrated that ELKS was significantly reduced by P7 and completely ablated by P9. Using confocal and electron microscopy we found that combined loss of CAST and ELKS resulted in significant reductions in calyx size. However, analysis of AZ ultrastructure revealed an increase in AZ size. In addition, we found a reduction in all Cav2 subtype currents and a reduction in Cav2.1 channel numbers. However, we found no change in synaptic transmission and short-term plasticity. Taken together our results implicate CAST/ELKS as key regulators of presynaptic morphology and active zone development. Furthermore, they regulate all presynaptic Cav2 subtype current levels and not just Cav2.1. Therefore, our results identify a role for CAST/ELKS proteins in the regulation of presynaptic maturation in a developing auditory brainstem circuit.

## Results

### Ablation of CAST in calyx of Held leads to increased levels of ELKS protein

To analyze the roles of CAST/ELKS we used the *Cast* KO/*Elks1*(CKO) transgenic mouse which CAST expression constitutively ablates and in the presence of Cre recombinase (Cre) ablates all of Elks isoforms (α, β, and γ) (Dong et al., 2018; Hagiwara et al., 2018). To do so, we injected our HdAd vector which independently co-expresses Cre and EGFP at P1. We chose the P1 time point, since the calyx axon at this time point has already innervated the Superior Olivary Complex (SOC) nuclei (Kandler and Friauf, 1993) but is still before the calyx terminal is established since it goes through period of extensive growth from P1 to P9 (Kandler and Friauf, 1993; Hoffpauir et al., 2006). Therefore, this enables to specifically look at calyx development and avoid potential phenotypes due to axon outgrowth or possible compensation by other signaling pathways early in development.

Although CAST/ELKS levels are characterized in the mature calyx, from P16 onwards, (Dong *et al.*, 2018), in the immature calyx of Held CAST/ELKS levels were unknown. Furthermore, it was unknown how early ELKS could be ablated *in vivo* as it is reported that in *in vitro* cultured neurons CAST/ELKS proteins have a half-life of ∼11-13 days (Fornasiero et al., 2018). Therefore, we first performed immunohistochemistry (IHC) using antibodies specific for CAST and ELKS, and VGLUT1 on immature, prehearing calyces (P9) in *Cast^+/+^/Elks^+/+^* and *Cast^−/−^/Elks^+/+^* terminals. P9 was chosen as time point since the calyx monoinnervates ∼90% of the principal cells of the MNTB (Holcomb et al., 2013). At P9 *Cast^+/+^/Elks^+/+^* calyces, we could only detect CAST not ELKS (Fig. 1A, B), while in *Cast^−/−^/Elks^+/+^* calyces, loss of CAST was accompanied by prominent expression of ELKS protein (Fig. 1C, D). Subsequently, IHC results from *Cast^−/−^/Elks^−/−^*calyces revealed that ELKS was undetectable by P9 (Fig. 1E, F). To determine how early ELKS could be completely ablated, we carried out analysis of ELKS levels at P7 *Cast^−/−^/Elks^−/−^*calyces. ROI of individual calyces were selected as VGLUT1 signal and same ROIs were applied to images from ELKS signal (Figure 2 A, B). P7 was chosen as time point when the calyx has started to reach a growth plateau (Holcomb et al., 2013). Results from our analysis revealed that at P7 ELKS levels were severely reduced in *Cast^−/−^ Elks ^−/−^*calyces (0.65 ± 0.12 *Cast^−/−^ Elks ^−/−^* vs 1.36 ± 0.17 *Cast^−/−^/Elks^+/+^*, n= 20 both groups, p<0.0001, t-test, Fig. 2C). In summary, our results in immature calyx of Held show that, CAST is the dominant isoform and ELKS levels are increased in the presynaptic terminal in the absence of CAST. Since, ELKS levels are dramatically reduced by P7 with undetectable levels by P9, this indicates that its half-life in the calyx of Held is significantly shorter than previous reports in other brain areas of 11-13 days (Fornasiero et al., 2018).

**Figure 1.**
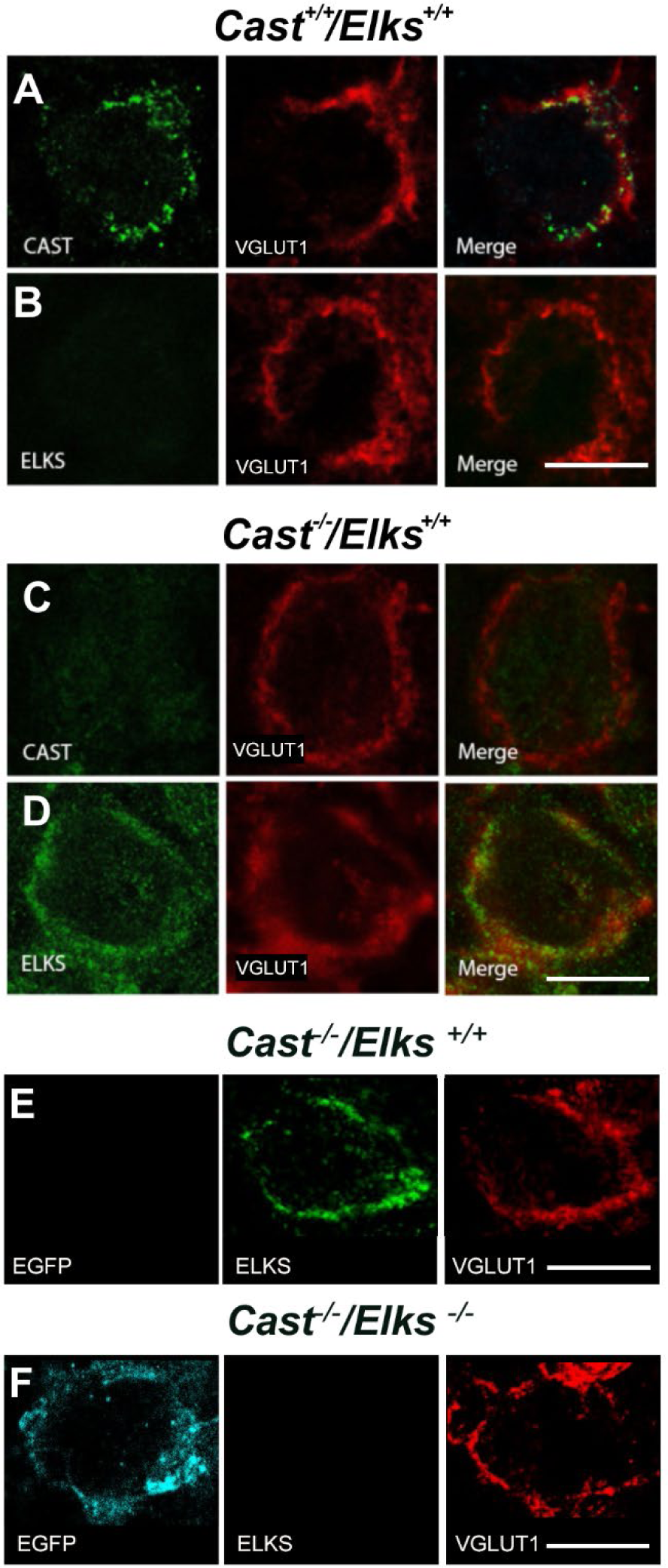
CAST is dominant isoform at immature calyx of Held, while both CAST and ELKS are not detectable by P9 in DKO animals. (A, C) Representative immunohistochemistry images showing CAST (green) and VGLUT1 (red) in *Cast^+/+^/Elks^+/+^* and *Cast^−/−^/Elks^+/+^* immature calyces (p9). (B, D) Immunohistochemistry showing ELKS proteins (green) and VGLUT1 (red) in *Cast^+/+^ /Elks^+/+^* and *Cast^−/−^ /Elks^+/+^* calyces. (E, F) EGFP (cyan) positive and negative calyces from *Cast^−/−^/Elks^fl/fl^* animals with ELKS (green) and VGLUT1 (red) staining, respectively. Scale bar 10 µm.

**Figure 2.**
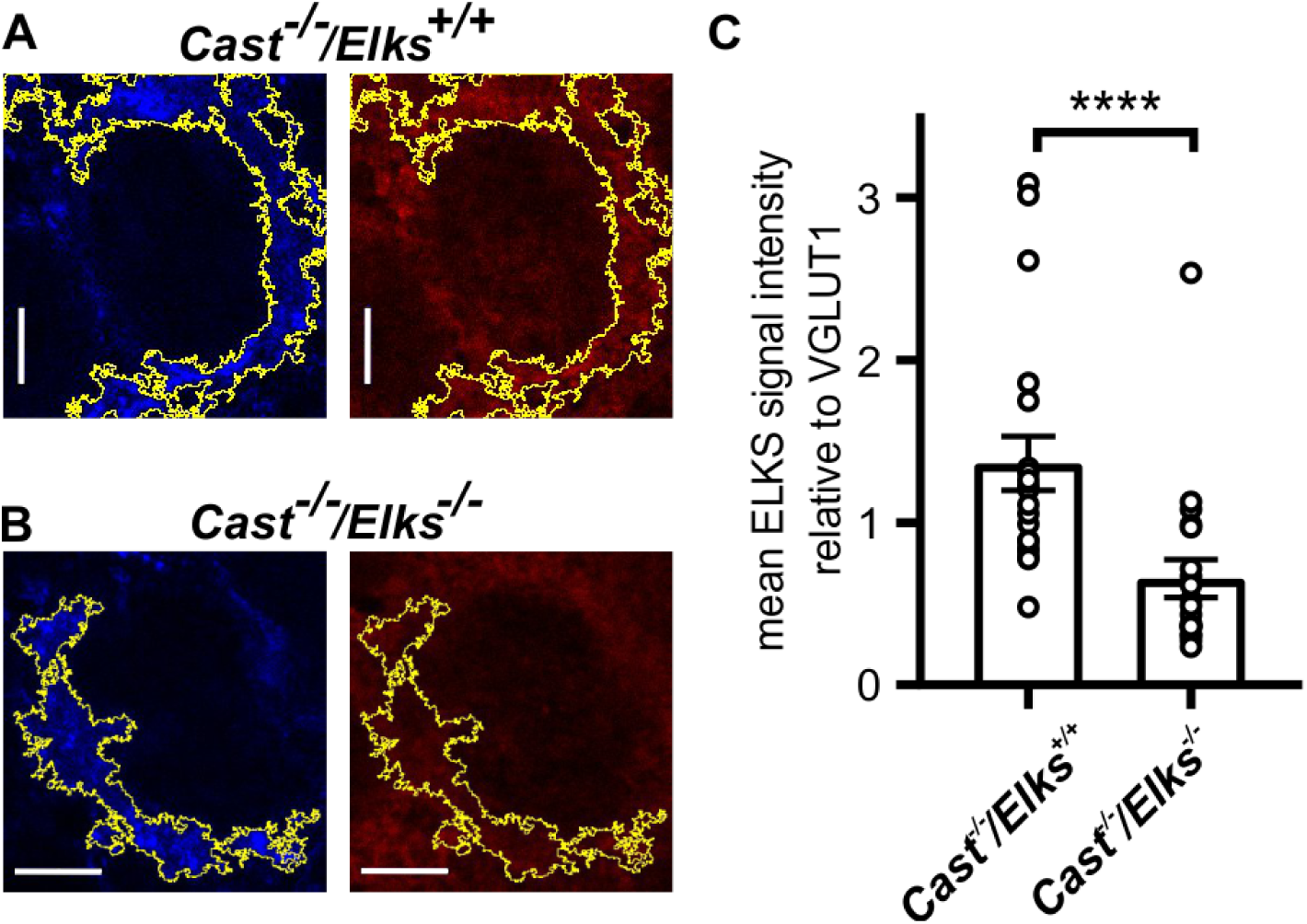
By P7 ELKS levels at the calyx are significantly reduced in DKO animals. (A) Immunohistochemical images of representative calyces of Held against VGLUT1 (blue) and ELKS (red) at *Cast^−/−^/Elks^+/+^* and (B) *Cast^−/−^ Elks ^−/^* animals, scale bar 5 µm. (C) Ratio of mean ELKS signal intensity relative to VGLUT1 signal. Data are represented as mean ± SEM. All groups n=20, t-test.

### Loss of CAST/ELKS impacts calyx size and active zone ultrastructure

Since we could significantly deplete ELKS levels by P7 with a complete loss of ELKS by P9, we set out to determine how loss of CAST/ELKS impacted calyx development. To do so, we carried out 3D reconstructions from confocal z-stacks images from *Cast^+/+^/Elks^+/+^*, *Cast^−/−^/Elks^+/+^*, *Cast^+/+^/Elks^−/−^*, *Cast^−/−^/Elks^−/−^* on mEGFP positive terminals of P9 calyces and determined the surface area and volume (Fig. 3). Analysis revealed reduction of surface area (1194 ± 48.9 *Cast^−/−^/Elks^+/+^* vs 1673 ± 60.6 µm^2^ *Cast^+/+^/Elks^+/+^*, n= 50, p<0.0001, Dunnett’s test) and volume (593.7 ± 25.2 *Cast^−/−^/Elks^+/+^* vs 714.6 ± 33.8 µm^3^ *Cast^+/+^/Elks^+/+^*, n= 50, p=0.02, Dunnett’s test) only in *Cast^−/−^/Elks^−/−^* compared to wild-type, while loss of either CAST or ELKS alone had no effect (Table 1).

**Figure 3.**
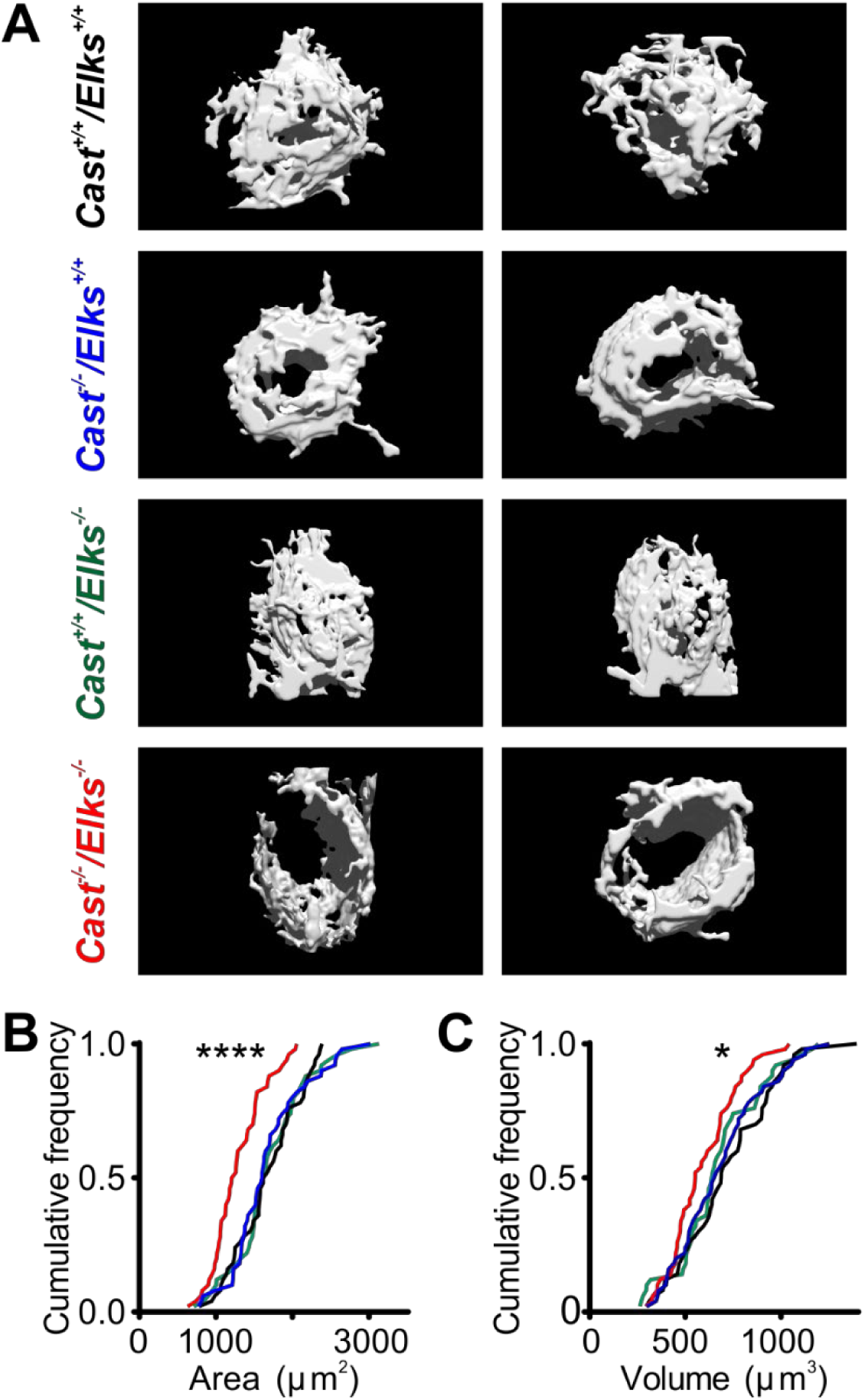
CAST/ELKS regulate volume and surface area of presynaptic terminals. (A) Representative reconstructions of calyx of Held terminals from *Cast^+/+^/Elks^+/+^*, *Cast^−/−^ /Elks^+/+^*, *Cast^+/+^/Elks^−/−^* and *Cast^−/−^/Elks^−/−^* P9 animals (two examples for each group). (B) Cumulative frequency presentation of population data for surface area and (C) volume of the presynaptic terminals. All groups n = 50.

**Table 1.**
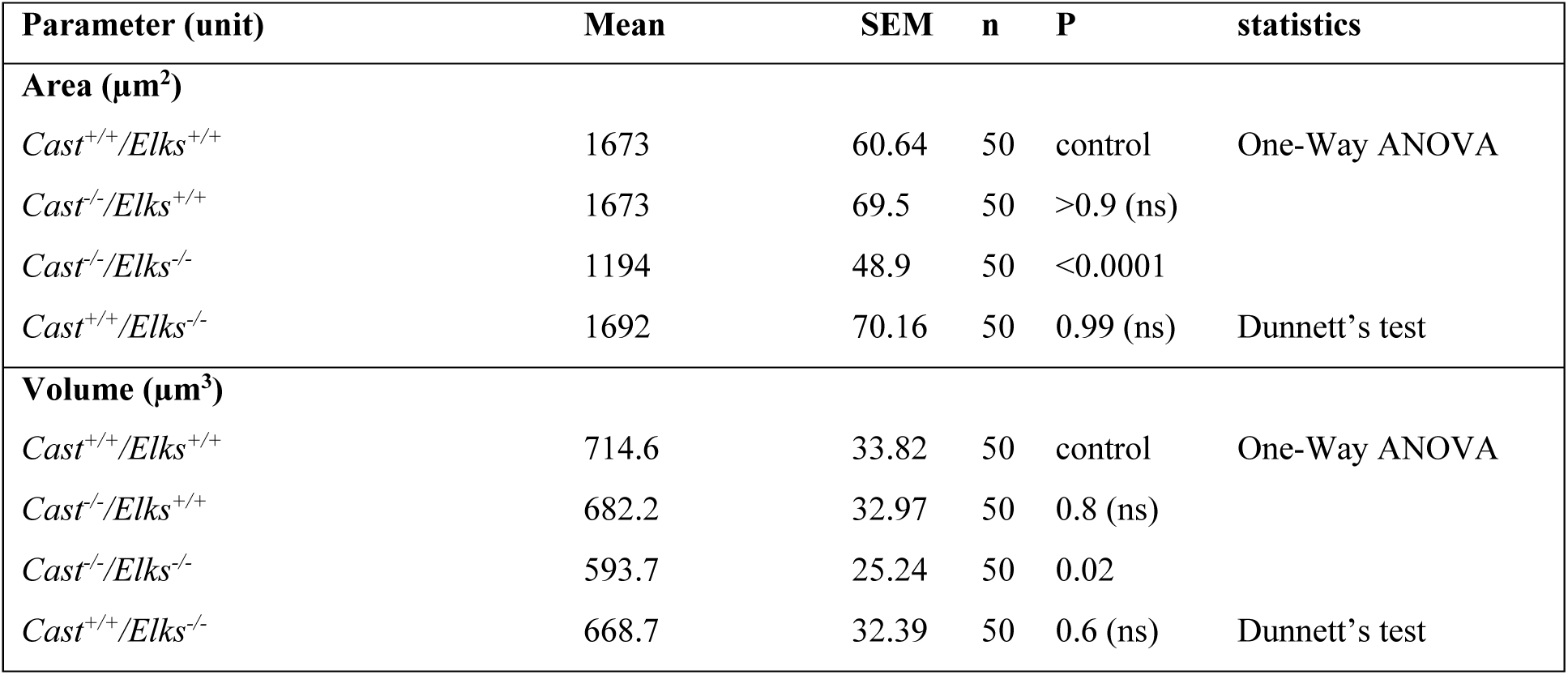
Analysis of Imaris reconstructions.

Although we identified a role for CAST/ELKS in regulating calyx morphology, it was important to determine how loss of CAST/ELKS impacted AZ ultrastructure. Therefore, we acquired and analyzed TEM images on ultrathin sections from *Cast^+/+^/Elks^+/+^, Cast^−/−^ /Elks^+/+^*, and *Cast^−/−^/Elks^−/−^* calyces to assess if there were alterations in AZ length, SV docking and distribution at P9-11 (Fig. 4, Table 2). Our analysis revealed a significant increase in AZ length only in the *Cast^−/−^/Elks^−/−^* calyces (Fig. 4 A, B) compared to *Cast^+/+^/Elks^+/+^* and *Cast^−/−^ /Elks^+/+^* (671.1 ± 27, n = 141, *Cast^−/−^ Elks ^−/−^*;466.3 ± 16.8, n = 126, *Cast^−/−^/Elks^+/+^*; 423.6 ± 13.02 µm, n= 124, *Cast^+/+^/Elks^+/+^*, p<0.0001, Kruskal-Wallis test). Subsequently, we measured SV docking, those within 5 nm (Taschenberger et al., 2002) of the membrane, and SV distribution. We found a slight increase in the docked SV numbers (Fig. 4D), in the *Cast^−/−^/Elks^+/+^, Cast^−/−^/Elks^−/−^* compared to *Cast^+/+^/Elks^+/+^* (3.1 ± 0.1, n = 141, *Cast^−/−^ Elks ^−/−^*; 3.0 ± 0.1, n = 126, *Cast^−/−^/Elks^+/+^*; 2.5 ± 0.1, n = 124, *Cast^+/+^/Elks^+/+^*, p=0.002, Kruskal-Wallis test; Fig. 4D, Table 2), but with no other changes in SV distribution. Taken together, this data demonstrates that in contrast to the mature calyx (Dong et al., 2018), deletion of CAST/ELKS results in a decrease in calyx size and an increase in AZ length. Therefore, we conclude that CAST/ELKS are involved in the regulation calyx morphology and AZ development during early calyx development. Furthermore, CAST and ELKS are functionally redundant in regulating the development of the calyx of Held.

**Figure 4.**
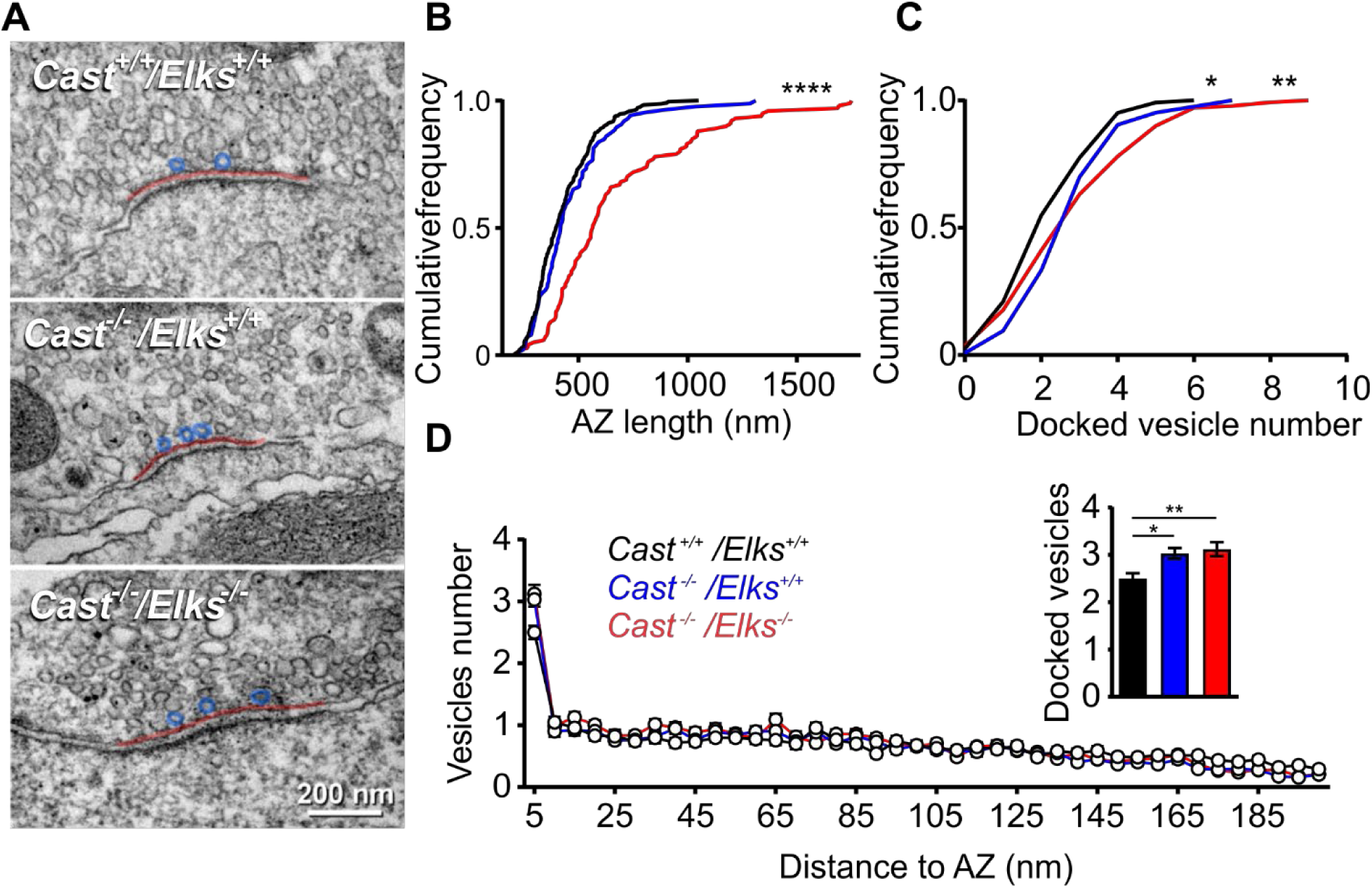
Loss of CAST/ELKS leads to increase in docked vesicle numbers and AZ length. (A) Representative electron micrographs of AZs taken from *Cast*^+/+^/*Elks*^+/+^ (top), *Cast^−/−^ Elks^+/+^* (middle), *Cast^−/−^Elks^−/−^* (bottom) where red line represents length of AZ and blue circles docked vesicles. (B, C) Cumulative frequency distribution of AZ length and docked vesicle numbers, respectively. (D) Distribution of docked vesicles as a function of their distance to AZ. Inset: mean population data. Data are represented as mean ± SEM.

**Table 2.** Ultrathin section Electron-microscopy data.

| Parameter (unit) | Mean [median] | SEM | n | P | statistics |
| --- | --- | --- | --- | --- | --- |
| **AZ length (nm)** |  |  |  |  |  |
| *Cast*<sup>+/+</sup>/*Elks*<sup>+/+</sup> | 423.6 [389.4] | 13 | 124 | control | Kruskal-Wallis |
| *Cast*<sup>-/-</sup>/*Elks*<sup>+/+</sup> | 466.3 [417.6] | 16.8 | 126 | ns |  |
| *Cast*<sup>-/-</sup>/*Elks*<sup>-/-</sup> | 671.1 [576.7] | 27 | 141 | p<0.0001 (****) | Dunn's test |
| **Number of docked SV** |  |  |  |  |  |
| *Cast*<sup>+/+</sup>/*Elks*<sup>+/+</sup> | 2.5 [2] | 0.1 |  | control | Kruskal-Wallis |
| *Cast*<sup>-/-</sup>/*Elks*<sup>+/+</sup> | 3.0 [3] | 0.1 |  | p=0.012 (*) |  |
| *Cast*<sup>-/-</sup>/*Elks*<sup>-/-</sup> | 3.1 [3] | 0.1 |  | p=0.002 (**) | Dunn's test |

\*AZ- active zone; SV- synaptic vesicle.

### Loss of CAST/ELKS results in a reduction of all Cav2 subtype currents

At the mature calyx which is Cav2.1 exclusive (Iwasaki and Takahashi, 1998), CAST/ELKS are positive regulators of presynaptic Cav2.1 channel density and Cav2 voltage-dependent activation (Dong *et al.*, 2018). However, it is unclear if CAST/ELKS regulation is specific to Cav2.1 or it affects all Cav2 subtypes. Unlike the mature calyx, the immature calyx is a mixed Cav2 subtype presynaptic terminal, Cav2.1 dominated but contains significant fraction of Cav2.2 and Cav2.3 currents (Doughty et al., 1998; Iwasaki and Takahashi, 1998). Therefore, it was important to determine whether CAST/ELKS regulation of Cav2 channels was specific to the developmental stage and Cav2.1 channel. To address this question, we carried out presynaptic whole-cell voltage-clamp recording at P9 calyces of *Cast^+/+^/Elks^+/+^*, *Cast^−/−^/Elks^+/+^* and *Cast^−/−^/Elks^−/−^* in the presence of calcium channel blockers specific to each Cav2 subtype (Fig. 5). Blockers were added in sequential order, ω-agatoxin IVA (for selective block of Cav2.1), followed by ω-conotoxin GVIA (for block of Cav2.2) and remaining calcium current (I_Ca_) was blocked using cadmium, a non-specific Ca^2+^ channel blocker. Our results revealed that presynaptic deletion of CAST/ELKS resulted in a reduction in total I_Ca_ (Fig. 5A, C) (0.56 ± 0.09 nA *Cast^−/−^ Elks ^−/−^*, 1.4 ± 0.11 nA *Cast^−/−^/Elks^+/+^*, 1.19 ± 0.14 nA *Cast^+/+^/Elks^+/+^*, p=0.0044, Dunnett’s test, Table 3). Analysis of the Cav2 subtype contributions revealed no change in the relative contributions (agatoxin: 69.88 ± 6.34 % *Cast^−/−^ Elks ^−/−^*, 66.96 ± 6.15 % *Cast^−/−^/Elks^+/+^*. 78.82 ± 6.9 %, *Cast^+/+^/Elks^+/+^*; conotoxin: 22.25 ± 8.2% *Cast^−/−^ Elks ^−/−^*, 18.49 ± 2.14% *Cast^−/−^/Elks^+/+^*, 12.1 ± 3.7 % *Cast^+/+^/Elks^+/+^*; cadmium: 7.9 ± 2.45 *Cast^−/−^ Elks ^−/−^*,14.55 ± 5.24% *Cast^−/−^/Elks^+/+^*, 9.1 ± 3.5 % *Cast^+/+^/Elks^+/+^*, n = 5 all groups, p=0.96; Dunnett’s test, Table 3) of each Cav2 subtype in the absence of CAST/ELKS (Fig. 5D).

**Figure 5.**
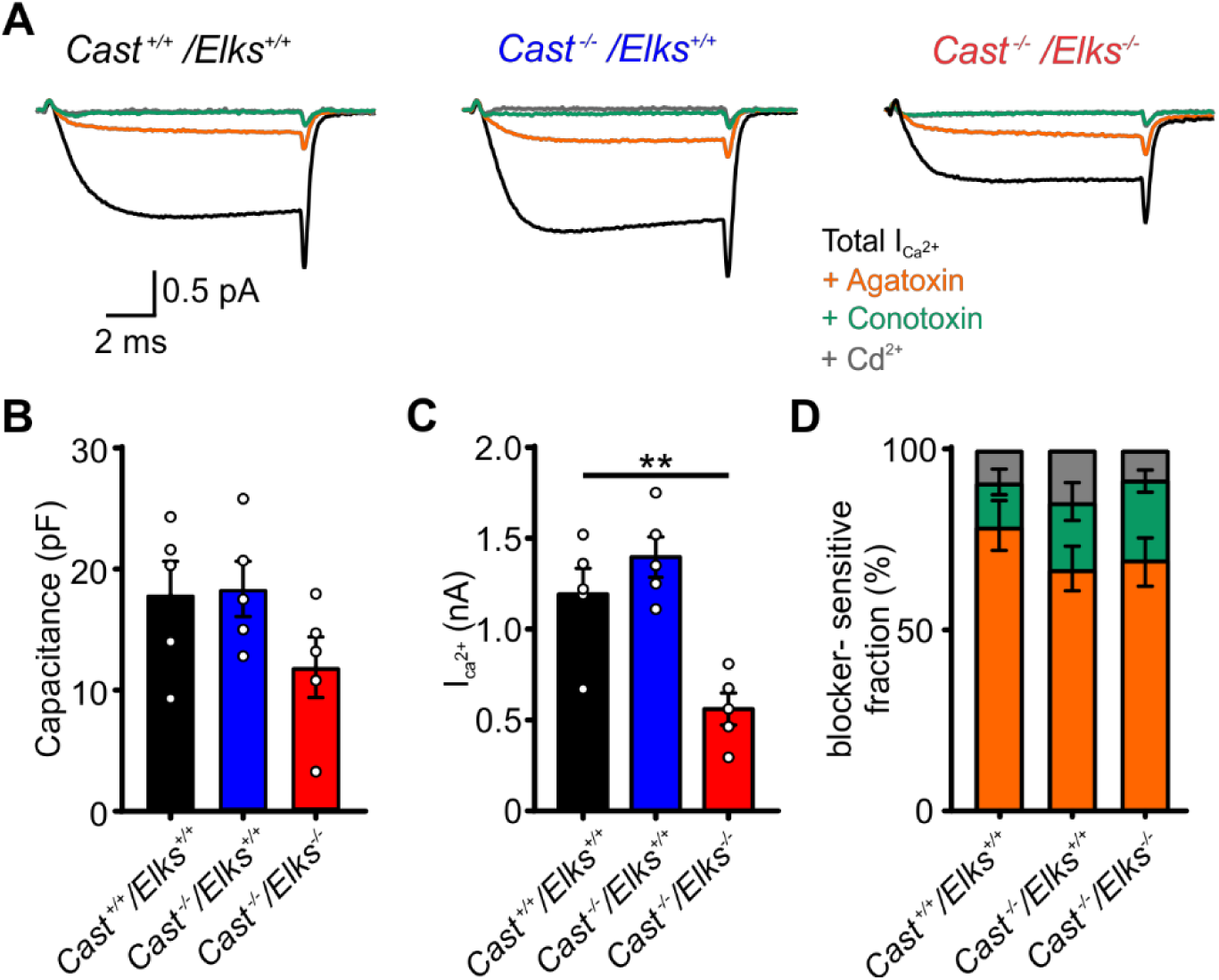
Deletion of CAST/ELKS affects all presynaptic Ca^2+^ channel subtypes. (A) Traces representing I_Ca_ subtypes from *Cast^+/+^/Elks^+/+^*, *Cast^−/−^/Elks^+/+^* and *Cast^−/−^/Elks^−/−^*. (B) Population data for cell capacitance. (C) Total calcium currents for all genotypes. (D) Relative fractions of different I_Ca_ subtypes (*Cast^+/+^/Elks^+/+^*, *Cast^−/−^/Elks^+/+^* n = 5, *Cast^−/−^/Elks^−/−^* n = 5, One Way ANOVA, p<0.01).

**Table 3:** Calcium channel subtype presynaptic recordings.

| Parameter (unit) | Mean/% of I<sub>Ca2+</sub> | SEM | n | P | statistics |
| --- | --- | --- | --- | --- | --- |
| **Total I<sub>Ca2+</sub> (nA)** |  |  |  |  |  |
| *Cast*<sup>+/+</sup>/*Elks*<sup>+/+</sup> | 1.19 | 0.14 | 5 | control | One-Way |
| *Cast*<sup>-/-</sup>/*Elks*<sup>+/+</sup> | 1.4 | 0.11 | 5 | 0.384 (ns) | ANOVA |
| *Cast*<sup>-/-</sup>/*Elks*<sup>-/-</sup> | 0.56 | 0.09 | 5 | 0.0044 (**) | Dunnett's test |
| **Agatoxin sensitive I<sub>Ca2+</sub> (nA)** |  |  |  |  |  |
| *Cast*<sup>+/+</sup>/*Elks*<sup>+/+</sup> | 0.91 / 78.82 | 0.09 | 5 | control | One-Way |
| *Cast*<sup>-/-</sup>/*Elks*<sup>+/+</sup> | 0.93 / 66.96 | 0.09 | 5 | 0.8 (ns)/0.35 | ANOVA |
| *Cast*<sup>-/-</sup>/*Elks*<sup>-/-</sup> | 0.38 / 69.88 | 0.05 | 5 | 0.012 (**)/ 0.53 | Dunnett's test |
| **Conotoxin sensitive I<sub>Ca2+</sub> (nA)** |  |  |  |  |  |
| *Cast*<sup>+/+</sup>/*Elks*<sup>+/+</sup> | 0.16 / 12.1 | 0.05 | 5 | control | One-Way |
| *Cast*<sup>-/-</sup>/*Elks*<sup>+/+</sup> | 0.26 / 18.49 | 0.05 | 5 | 0.36 (ns)/0.62 | ANOVA |
| *Cast*<sup>-/-</sup>/*Elks*<sup>-/-</sup> | 0.14 / 22.25 | 0.05 | 5 | 0.82 (ns)/0.33 | Dunnett's test |
| **Cd<sup>2+</sup> sensitive I<sub>Ca2+</sub> (nA)** |  |  |  |  |  |
| *Cast*<sup>+/+</sup>/*Elks*<sup>+/+</sup> | 0.12 / 9.1 | 0.06 | 5 | control | One-Way |
| *Cast*<sup>-/-</sup>/*Elks*<sup>+/+</sup> | 0.2 / 14.55 | 0.06 | 5 | 0.46 (ns)/0.53 | ANOVA |
| *Cast*<sup>-/-</sup>/*Elks*<sup>-/-</sup> | 0.04 / 7.87 | 0.01 | 5 | 0.46 (ns)/0.96 | Dunnett's test |

Since I_Ca_ voltage-dependence of activation changes with calyx development (Chen et al., 2015), we measured the current-voltage (IV) relationships at P9-11 calyces using presynaptic whole-cell voltage-clamp recordings in the various genotypes. In this case, we switched to 1 mM external Ca^2+^ to minimize the potential influence of voltage-clamp errors. Analysis of the I_Ca_ current-voltage relationship revealed a reduction in the maximum I_ca_ steady state current in *Cast^−/−^Elks^−/−^* calyces compared to *Cast^+/+^ Elks^+/+^* and *Cast^−/−^ Elks^+/+^* (525 ± 40 pA, n= 12, *Cast^−/−^ Elks ^−/−^*; 903 ± 53 pA, n = 10, *Cast^+/+^/Elks^+/+^*; 867 ± 69 pA, n = 10, *Cast^−/−^/Elks^+/+^*, p = 0.0001, Dunnett’s test,; Fig. 6C, Table 4). Fits according to Hodgkin-Huxley formalism of the individual IV curves revealed a small change in voltage dependence of activation (8.43 ± 0.4 mV, n= 12, *Cast^−/−^ Elks ^−/−^*; 6.81± 0.5 mV, n = 10, *Cast^−/−^/Elks^+/+^* 6.64 ± 0.4 mV, n =10, *Cast^+/+^/Elks^+/+^*; p=0.04, Dunn’s test, Table 4) in *Cast^−/−^Elks^−/−^*(Fig. 6D). Analysis of the tail currents confirmed a reduction in I_Ca_, but no significant change in voltage-dependence of activation was detected using Boltzmann fitting function although a tendency to a small rightward shift was seen (8.6 ± 0.6 mV, n = 12, *Cast^−/−^ Elks ^−/−^*; 7.7 ± 0.7 mV, n = 10, *Cast^−/−^/Elks^+/+^*; 8.7 ± 0.4 mV, n = 10, *Cast^+/+^/Elks^+/+^*, p = 0.99, Dunnett’s test; Fig. 6E, F; Table 4). Based on these results, we conclude that CAST/ELKS is a critical regulator of presynaptic Cav2 subtype levels and is not specific to Cav2.1.

**Figure 6.**
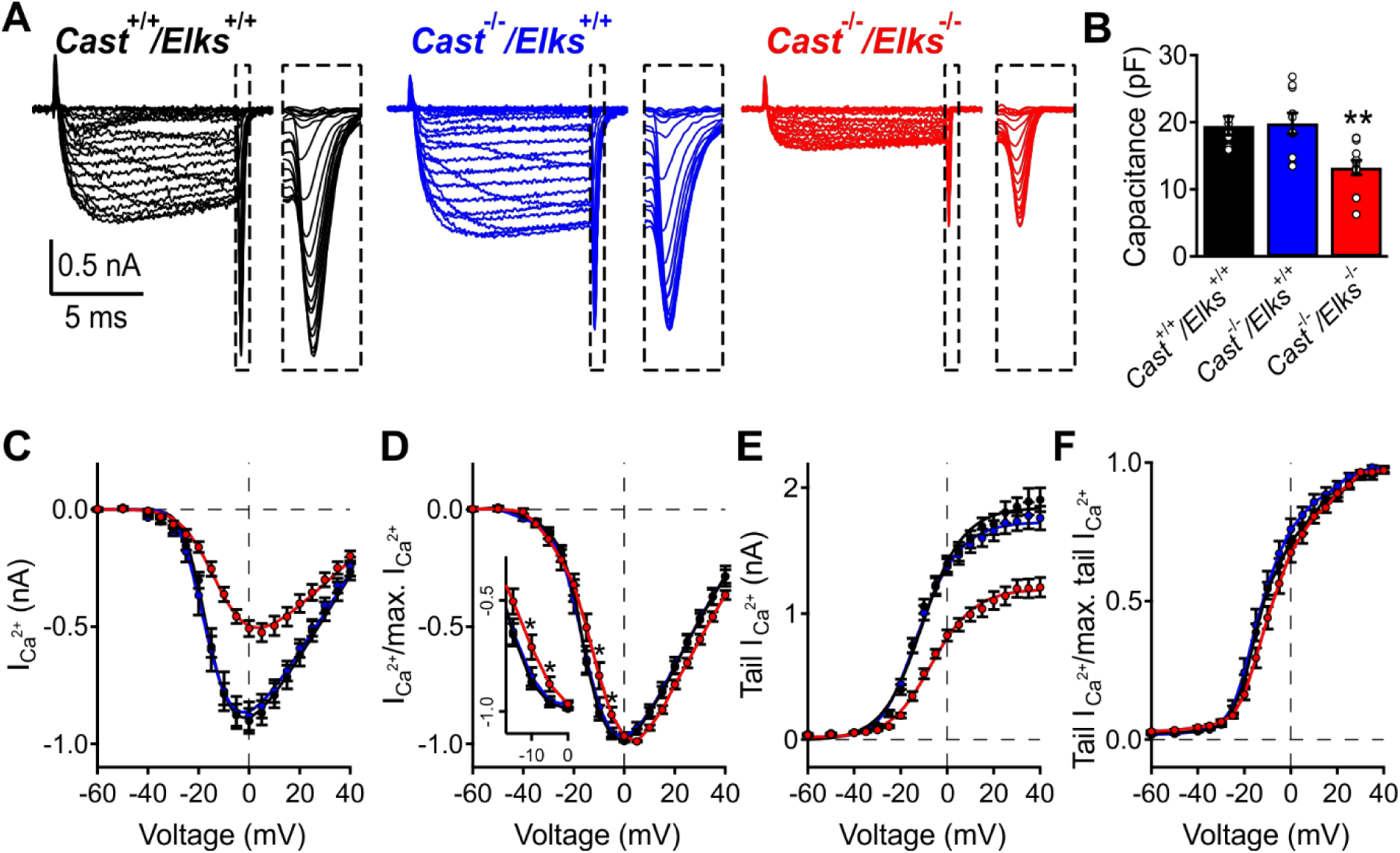
Ablation of CAST/ELKS leads to reduction in I_Ca_ and rightward shift in the voltage-dependence of activation of presynaptic Cav2 channels. (A) Representative traces of presynaptic recordings of I_Ca_ from *Cast^+/+^ Elks^+/+^* (black), *Cast^−/−^ Elks^+/+^* (blue), *Cast^−/−^ Elks^−/−^* (red). Inset depicts enlarged tail currents. (B) Population data for capacitance for all tested groups. (C) Mean peak Ca^2+^ currents as a function of voltage. (D) Mean Ca^2+^ normalized to the maximum peak as function of voltage. Inset: voltage range −15 – 0 mV. (E, F) Absolute values of peak Ca^2+^ tail currents and normalized to the maximum tail current as a function of voltage, respectively. Data are represented as mean ± SEM, *Cast^+/+^ Elks^+/+^* n= 10, *Cast^−/−^ Elks^+/+^* n=10, *Cast^−/−^Elks^−/−^* n= 12, *p<0.05, **p<0.01, One-way ANOVA.

**Table 4.** Presynaptic recordings summary values.

| Parameter (unit) | Mean | SEM | n | P | statistics |
| --- | --- | --- | --- | --- | --- |
| **max. I<sub>Ca</sub> (pA)** |  |  |  |  |  |
| *Cast*<sup>+/+</sup>/*Elks*<sup>+/+</sup> | 903 | 53 | 10 | control | One Way |
| *Cast*<sup>-/-</sup>/*Elks*<sup>+/+</sup> | 867 | 69 | 10 | 0.848 (ns) | ANOVA |
| *Cast*<sup>-/-</sup>/*Elks*<sup>-/-</sup> | 525 | 40 | 12 | 0.0001 (***) | Dunnett's test |
| **Membrane capacitance C<sub>slow</sub> (pF)** |  |  |  |  |  |
| *Cast*<sup>+/+</sup>/*Elks*<sup>+/+</sup> | 19.5 | 1.4 | 10 | control | Kruskal-Wallis |
| *Cast*<sup>-/-</sup>/*Elks*<sup>+/+</sup> | 19.8 | 1.6 | 10 | >0.999 |  |
| *Cast*<sup>-/-</sup>/*Elks*<sup>-/-</sup> | 13.3 | 1.1 | 12 | 0.0046 (**) | Dunn's test |
| **Ca<sup>2+</sup> currents as function of voltage** |  |  |  |  |  |
| **Half-maximal activation voltage V<sub>m</sub> (mV)** |  |  |  |  |  |
| *Cast*<sup>+/+</sup>/*Elks*<sup>+/+</sup> | -23.6 | 0.8 | 10 | control | One Way |
| *Cast*<sup>-/-</sup>/*Elks*<sup>+/+</sup> | -24.1 | 1.0 | 10 | 0.9516 (ns) | ANOVA |
| *Cast*<sup>-/-</sup>/*Elks*<sup>-/-</sup> | -23.3 | 1.7 | 12 | 0.983 (ns) | Dunnett's test |
| **Voltage-dependence of activation k<sub>m</sub> (mV)** |  |  |  |  |  |
| *Cast*<sup>+/+</sup>/*Elks*<sup>+/+</sup> | 6.64 | 0.4 | 10 | control | Kruskal-Wallis |
| *Cast*<sup>-/-</sup>/*Elks*<sup>+/+</sup> | 6.81 | 0.5 | 10 | >0.999 (ns) |  |
| *Cast*<sup>-/-</sup>/*Elks*<sup>-/-</sup> | 8.43 | 0.4 | 12 | 0.04 (*) | Dunn's test |
| **Ca<sup>2+</sup> tail currents as function of voltage** |  |  |  |  |  |
| **Half-maximal activation voltage V<sub>0.5</sub> (mV)** |  |  |  |  |  |
| *Cast*<sup>+/+</sup>/*Elks*<sup>+/+</sup> | -10.6 | 1.0 | 10 | control | One Way |
| *Cast*<sup>-/-</sup>/*Elks*<sup>+/+</sup> | -11.4 | 1.9 | 10 | 0.8884 (ns) | ANOVA |
| *Cast*<sup>-/-</sup>/*Elks*<sup>-/-</sup> | -6.7 | 1.4 | 12 | 0.1242 (ns) | Dunnett's test |
| **Voltage-dependence of activation k (mV)** |  |  |  |  |  |
| *Cast*<sup>+/+</sup>/*Elks*<sup>+/+</sup> | 8.7 | 0.4 | 10 | control | One Way |
| *Cast*<sup>-/-</sup>/*Elks*<sup>+/+</sup> | 7.7 | 0.7 | 10 | 0.4207 (ns) | ANOVA |
| *Cast*<sup>-/-</sup>/*Elks*<sup>-/-</sup> | 8.6 | 0.6 | 12 | 0.9958 (ns) | Dunnett's test |

### Loss of CAST/ELKS results in decreased Cav2.1 numbers

Our data revealed that the reduction in I_Ca_ observed in the absence of CAST/ELKS was accompanied by a reduction in calyx size (Fig. 3B, C). However, it was unclear if the reduction of Cav2 currents was due to a reduction of calcium channel numbers in the calyx active zone or due to smaller calyces having less Cav2 channels. To resolve this question. we carried out SDS-FFRIL in combination with immunogold TEM on P9 calyces of *Cast^−/−^/Elks^+/+^* and *Cast^−/−^/Elks^−/−^* animals. *Cast^−/−^/Elks^+/+^* was used as a control because there were no changes in Ca^2+^ currents compared to *Cast^+/+^/Elks^+/+^* and served as an in-slice control to minimize potential sample-to-sample variability. We used Cav2.1-specific antibody (KO verified) in combination with an EGFP antibody that allowed for detection of transduced calyces. We then quantified the number of Cav2.1 channels by counting the number of 12 nm gold particles linked to the secondary antibody in both *Cast^−/−^/Elks^+/+^* and *Cast^−/−^/Elks^−/−^* calyces on the presynaptic P face that was in contact with the postsynaptic membrane. Clusters of Cav2.1 channels were defined as at least two gold particles within a 30 nm radius to account for labeling uncertainty (Althof et al., 2015; Dong et al., 2018). To determine how Cav2.1 density was affected in the calyx we measured the number of gold particles per cluster, cluster area, cluster particle density, number of clusters and total number of particles (Fig. 7, Table 5). Analysis revealed that loss of CAST/ELKS resulted in a mean reduction in gold particles/cluster (4.4 ± 0.26 vs 5.8 ± 0.43, p<0.05, Mann-Whitney U test) as well as a reduction in cluster area (0.0081 ± 0.0004 vs 0.0098 ± 0.0006 µm^2,^ p<0.05, Mann-Whitney U test) in *Cast^−/−^/Elks^−/−^* compared to *Cast^−/−^/Elks^+/+^* (Fig. 7A-C), and a reduction in gold particle density per cluster (520.0 ± 6.9 vs 556.4 ± 8.7 particles/µm^2^, p<0.01, Mann-Whitney U test; Fig. 7D). To determine if the loss of Cav2.1 channels is associated with a reduction in cluster number, we measured the total number of particles and analyzed the number of single particle population compared to the number of clusters. Our results revealed no change in the relative frequency of single gold particles (39% vs 43%; Fig. 7E) between the *Cast^−/−^/Elks^+/+^* and *Cast^−/−^/Elks^−/−^*.

**Figure 7.**
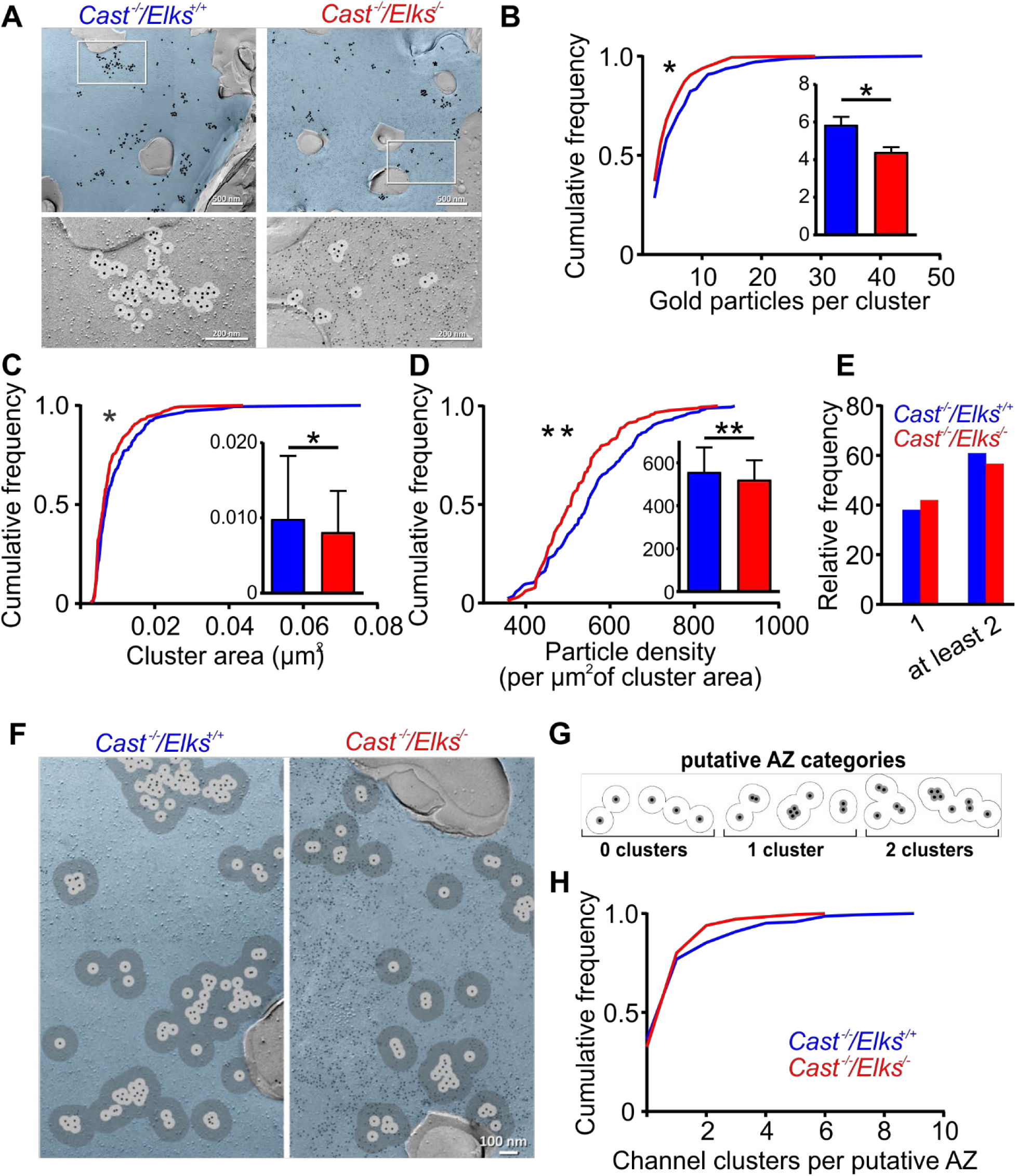
CAST/ELKS proteins ablation in the calyx of Held presynaptic terminal leads to reduction of Cav2.1 numbers and clusters at the AZ. (A) Representative SDS-FFRIL immuno-gold labeled replicas of calyx P-faces (pseudocolored blue) of *Cast^−/−^ Elks^+/+^* and *Cast^−/−^Elks^−/−^*. Top: Cav2.1 distribution labeled with large gold particles (12 nm) and small gold particles (6 nm) label mEGFP (specific for *Cast^−/−^Elks^−/−^* only). Bottom: High magnification images of the boxed areas on the top image. (B) Cumulative frequency distribution of gold particles per cluster, inset shows data points for *Cast^−/−^ Elks^+/+^* and *Cast^−/−^/Elks^−/−^*. (C) Cumulative frequency distribution of population data for cluster area. Inset: population data for *Cast^−/−^ Elks^+/+^* and *Cast^−/−^Elks^−/−^*. (D) Mean cumulative frequency distribution of number of particles per µm^2^, inset population data. (E) Percentage of single and at least two particles in the cluster. (F) SDS-FFRIL representative immuno-gold labeled replicas of putative AZ, where dark grey circles depict 100 nm radius from the cluster of gold particle labeled Cav2.1 channels and light grey circles depict 30 nm radius from Cav2.1 clusters. (G) Schematic presentation of putative AZ. (H) Cumulative frequency distribution of channel clusters per AZ. Data are represented as mean ± SEM, *p<0.05, **p<0.01, Mann-Whitney U.

**Table 5.**
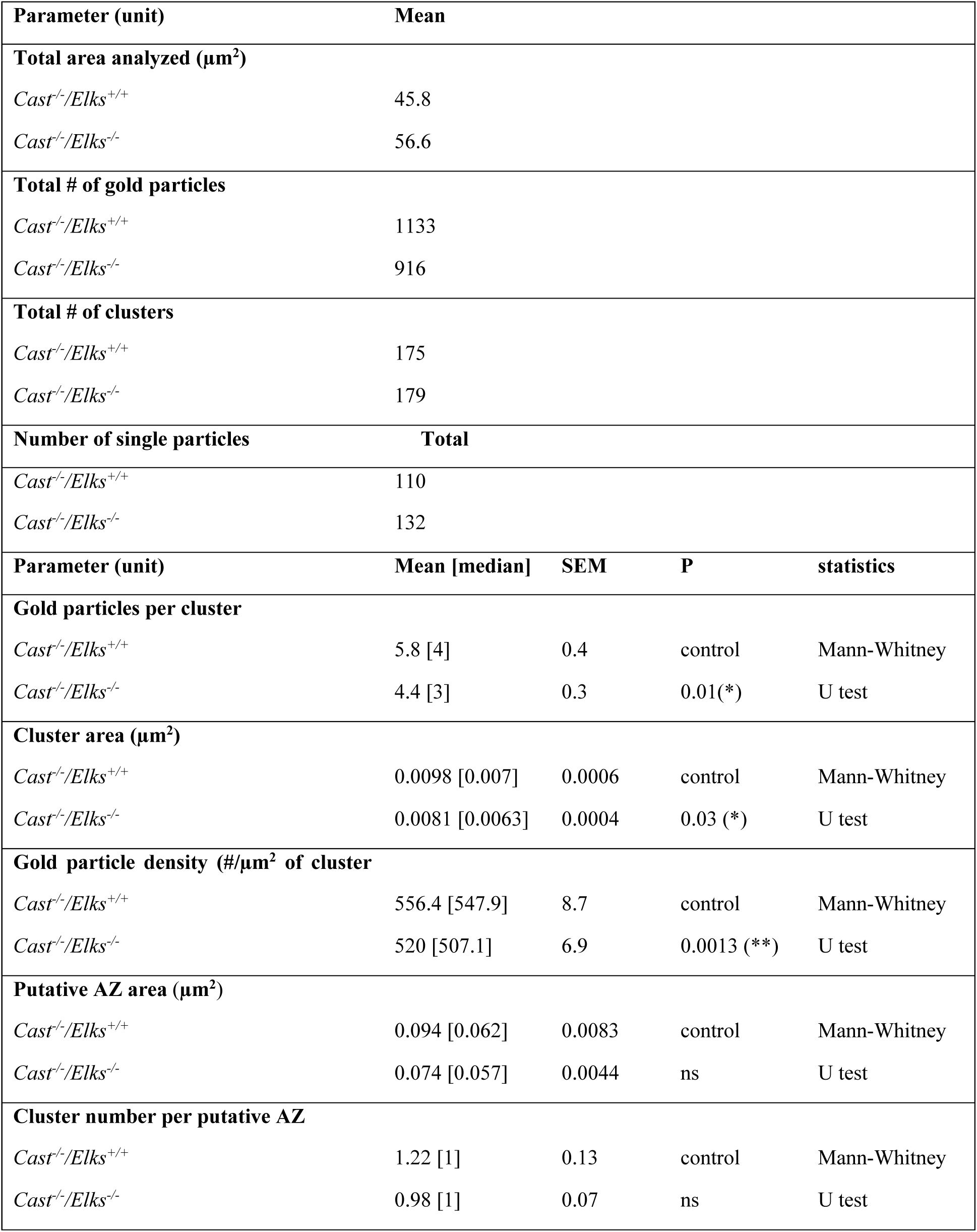
SDS-FFRIL summary values.

AZ can contain multiple clusters of Cav2 channels (Holderith et al., 2012; Lenkey et al., 2015; Maschi and Klyachko, 2017; Miki et al., 2017), thus it became important to know if there was a change in the number of clusters within the AZ at the calyx. Using a 100 nm radius around individual calcium channels to measure cluster area can be used as an approximate measure of AZ size at the calyx (Dong et al., 2018). Thus, we counted the number of Cav2.1-containing clusters within the AZ area approximated by using a 100 nm radius (Fig. 7F-H, Table 5). Single particles within the AZ area were not counted as cluster. We found that there was no difference in the AZ area and number of Cav2.1-containing clusters within individual AZs in the *Cast^−/−^/Elks^+/+^* and *Cast^−/−^/Elks^−/−^* calyces. Taken together, our results revealed that the reductions in Cav2 currents were correlated with a decrease in the Cav2.1 channel numbers in the AZ. Since, Cav2 currents were reduced in the absence of CAST/ELKS, we conclude that CAST/ELKS regulate Cav2 channel number and density in the prehearing calyx.

### Loss of CAST/ELKS does not impact AP evoked release, RRP size, release kinetics or spontaneous release

We identified that roles for CAST/ELKS in regulating calyx morphology, AZ ultrastructure and Cav2 levels in the immature calyx are distinct from the mature calyx. CAST/ELKS regulate *Pr*, RRP size and short-term plasticity of the mature calyx (Dong et al., 2018). Mature calyx that uses nanodomain release in contrast to the immature that utilizes microdomain release (Borst and Sakmann, 1996; Fedchyshyn and Wang, 2005). Therefore, we wanted to determine how loss of CAST/ELKS impacted synaptic transmission in the immature calyx and determine if this led to a similar synaptic transmission phenotype as in the mature calyx. To do so, we performed midline stimulation of calyx axons and recorded the AMPA receptors mediated excitatory postsynaptic currents (EPSCs) in whole cell voltage clamp mode from the principal cells of the MNTB, innervated by transduced or non-transduced calyces using *Cast^+/+^/Elks^+/+^*, *Cast^−/−^/Elks^+/+^* and *Cast^−/−^/Elks^−/−^* mice. We used a 0.05 Hz stimulation in 2 mM Ca^2+^ to measure basal AP-evoked synaptic transmission. We found that loss of CAST/ELKS had no effect on basal AP-evoked synaptic transmission (Table 6). Therefore, we next test if the loss of CAST/ELKS impacted *Pr* and the RRP at this developmental stage. We carried out afferent fiber stimulation at 100 Hz and determined the RRP size with the back-extrapolation method (Fig.8E) (Neher, 2015). Subsequent analysis revealed no change in RRP size (Fig.8G, Table 6) or SV replenishment rates (Fig. 8H, Table 6). In addition, there was no change in the release probability (Table 6) or paired pulse ratio (PPR) (Fig. 8I, Table 6).

**Figure 8.**
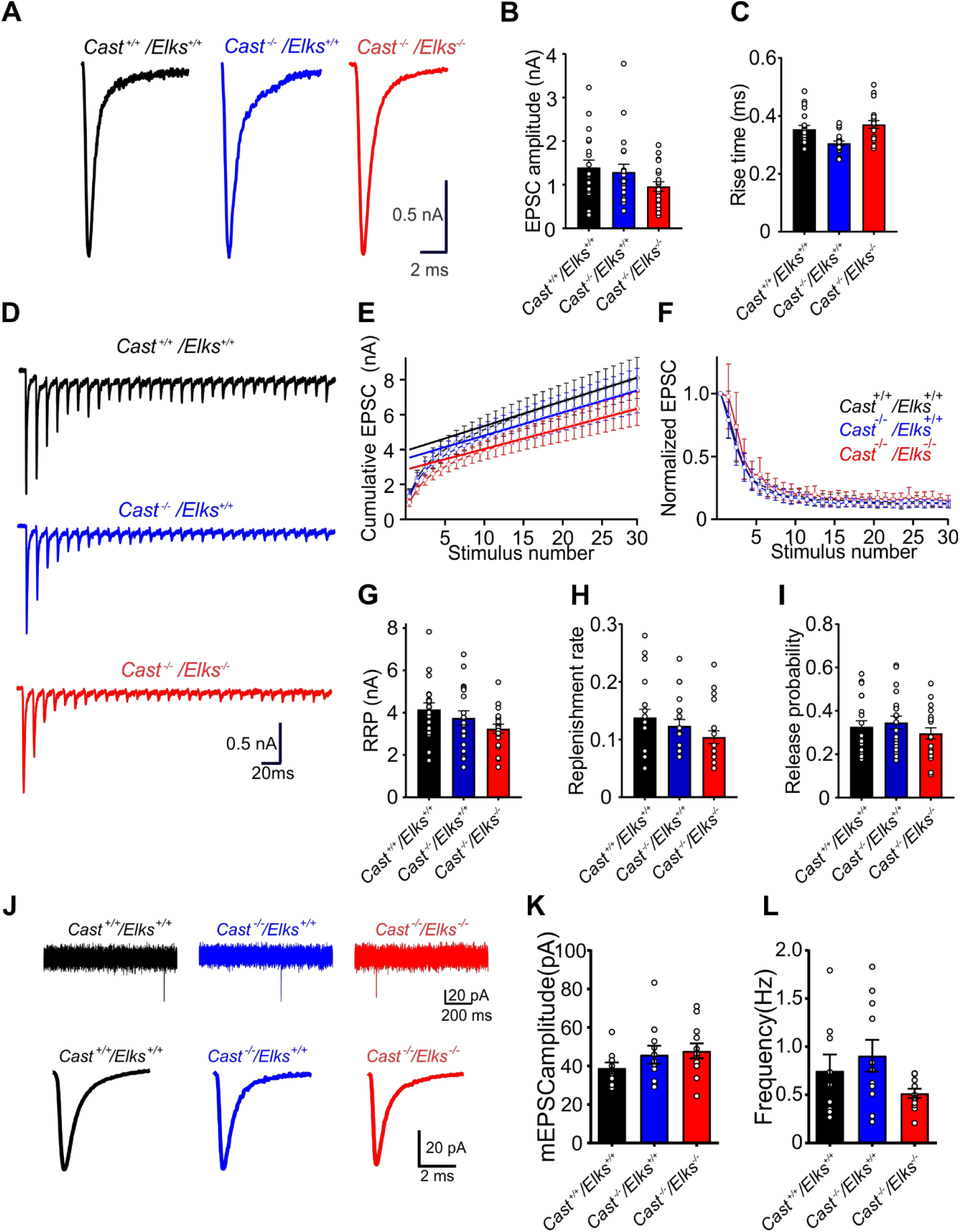
Action potential-evoked and spontaneous release is not impaired with loss of CAST and ELKS. (A) Representative recordings of single EPSC from *Cast^+/+^ Elks^+/+^*, *Cast^−/−^ Elks^+/+^*, *Cast^−/−^Elks^−/−^*. (B, C) Absolute EPSC amplitudes and rise times for all tested genotypes. (D) Representative recordings of AP evoked responses. (E) Cumulative EPSC amplitudes as a function of stimulus number with steady state back extrapolation line. (F) EPSC in the train normalized to the amplitude of the first peak. (G, H, I) Mean RRP size, replenishment rate, vesicular release probability for *Cast^+/+^ Elks^+/+^*, *Cast^−/−^ Elks^+/+^*, *Cast^−/−^ Elks^−/−^*. (J) Representative recordings of mEPSC for all genotypes, with average mEPSC waveform from the same cell, respectively. (K, L) Population data for mEPSC amplitude and frequency. Data are represented as a mean ± SEM, *Cast^+/+^ Elks^+/+^* n = 20, *Cast^−/−^ Elks^+/+^* n = 20, *Cast^−/−^Elks^−/−^* n = 20; mEPSCs: *Cast^+/+^ Elks^+/+^* n = 12, *Cast^−/−^ Elks^+/+^* n = 11, *Cast^−/−^Elks^−/−^* n = 12, *p<0.05, **p<0.01, ***p<0.001, One Way ANOVA.

**Table 6.** Afferent fiber stimulation data.

| Parameter (unit) | Mean | SEM | n | P | statistics |
| --- | --- | --- | --- | --- | --- |
| **Basal EPSC amplitude** |  |  |  |  |  |
| *Cast*<sup>+/+</sup>/*Elks*<sup>+/+</sup> | 1.37 | 0.16 | 20 | control | One-Way |
| *Cast*<sup>-/-</sup>/*Elks*<sup>+/+</sup> | 1.4 | 0.18 | 20 | >0.9999(ns) | ANOVA |
| *Cast*<sup>-/-</sup>/*Elks*<sup>-/-</sup> | 1.04 | 0.10 | 20 | 0.48 (ns) | Dunnett's test |
| **RRP (nA)** |  |  |  |  |  |
| *Cast*<sup>+/+</sup>/*Elks*<sup>+/+</sup> | 4.16 | 0.31 | 20 | control | Kruskal-Wallis |
| *Cast*<sup>-/-</sup>/*Elks*<sup>+/+</sup> | 3.75 | 0.34 | 20 | 0.512 (ns) |  |
| *Cast*<sup>-/-</sup>/*Elks*<sup>-/-</sup> | 3.24 | 0.21 | 20 | 0.06 (ns) | Dunn's test |
| **Replenishment rate** |  |  |  |  |  |
| *Cast*<sup>+/+</sup>/*Elks*<sup>+/+</sup> | 0.14 | 0.014 | 20 | control | Kruskal-Wallis |
| *Cast*<sup>-/-</sup>/*Elks*<sup>+/+</sup> | 0.12 | 0.011 | 20 | >0.9999(ns) |  |
| *Cast*<sup>-/-</sup>/*Elks*<sup>-/-</sup> | 0.10 | 0.011 | 20 | 0.094 (ns) | Dunn's test |
| **PPR** |  |  |  |  |  |
| *Cast*<sup>+/+</sup>/*Elks*<sup>+/+</sup> | 0.83 | 0.09 | 20 | control | Kruskal-Wallis |
| *Cast*<sup>-/-</sup>/*Elks*<sup>+/+</sup> | 0.8 | 0.07 | 20 | >0.9999(ns) |  |
| *Cast*<sup>-/-</sup>/*Elks*<sup>-/-</sup> | 0.98 | 0.12 | 20 | >0.9999(ns) | Dunn's test |
| **Pr** |  |  |  |  |  |
| *Cast*<sup>+/+</sup>/*Elks*<sup>+/+</sup> | 0.33 | 0.03 | 20 | control | One-Way |
| *Cast*<sup>-/-</sup>/*Elks*<sup>+/+</sup> | 0.35 | 0.3 | 20 | >0.9999(ns) | ANOVA |
| *Cast*<sup>-/-</sup>/*Elks*<sup>-/-</sup> | 0.3 | 0.02 | 20 | >0.9999(ns) | Dunnett's test |

\*RRP-readily releasable pool, PPR- paired pulse ratio, Pr-release probability, EPSC-excitatory postsynaptic current.

Spontaneous release can be regulated independently from evoked release mechanisms (Kavalali, 2014) and loss of CAST/ELKS results in increased spontaneous release rates in the mature calyx (Dong et al., 2018). Therefore, we set out to determine if deletion of CAST/ELKS resulted in a similar change in the P9-11 calyx. To do so, we measured miniature EPSC (mEPSC) frequency and amplitude. Our results revealed that the loss of CAST/ELKS did not cause a change in the mEPSC frequency (Table 7) or amplitude (Fig.8J-L, Table 7). Taken together, these data demonstrate that loss of CAST/ELKS does not impact AP-evoked or spontaneous release.

**Table 7.** Miniature excitatory postsynaptic current (mEPSC) recordings.

| Parameter (unit) | Mean | SEM | n | P | statistics |
| --- | --- | --- | --- | --- | --- |
| **Frequency (Hz)** |  |  |  |  |  |
| *Cast*<sup>+/+</sup>/*Elks*<sup>+/+</sup> | 0.78 | 0.12 | 12 | control | One-Way ANOVA |
| *Cast*<sup>-/-</sup> *Elks*<sup>+/+</sup> | 0.9 | 0.16 | 11 | 0.47 |  |
| *Cast*<sup>-/-</sup>/*Elks*<sup>-/-</sup> | 0.5 | 0.05 | 12 | 0.23 (ns) | Dunnett's test |
| **Amplitude (pA)** |  |  |  |  |  |
| *Cast*<sup>+/+</sup>/*Elks*<sup>+/+</sup> | 38.95 | 2.3 | 12 | control | One-Way ANOVA |
| *Cast*<sup>-/-</sup> *Elks*<sup>+/+</sup> | 45.81 | 4.7 | 11 | 0.47 (ns) |  |
| *Cast*<sup>-/-</sup>/*Elks*<sup>-/-</sup> | 47.82 | 3.9 | 12 | 0.12 (ns) | Dunnett's test |
| **Rise time (ms)** |  |  |  |  |  |
| *Cast*<sup>+/+</sup>/*Elks*<sup>+/+</sup> | 0.43 | 0.05 | 12 | control | Kruskal-Wallis |
| *Cast*<sup>-/-</sup> *Elks*<sup>+/+</sup> | 0.34 | 0.05 | 11 | 0.58 (ns) |  |
| *Cast*<sup>-/-</sup>/*Elks*<sup>-/-</sup> | 0.5 | 0.08 | 12 | 0.58 (ns) | Dunn's test |
| **Half-width (ms)** |  |  |  |  |  |
| *Cast*<sup>+/+</sup>/*Elks*<sup>+/+</sup> | 1.34 | 0.12 | 12 | control | One-Way ANOVA |
| *Cast*<sup>-/-</sup> *Elks*<sup>+/+</sup> | 0.99 | 0.11 | 11 | 0.34 |  |
| *Cast*<sup>-/-</sup>/*Elks*<sup>-/-</sup> | 1.51 | 0.28 | 12 | 0.75 (ns) | Dunnett's test |

### Loss of CAST/ELKS does not change the RRP or total releasable pool release kinetics

Since the loss of CAST/ELKS resulted in morphological changes but no changes in synaptic transmission it was possible that the lack of effects on synaptic transmission were due to tighter SV to Cav2 coupling, a shift from microdomain to nanodomain (Stanley, 2016). To determine if there were changes in coupling, we performed paired whole-cell voltage-clamp recordings of the pre and postsynaptic compartment of at P9-11the calyx/MNTB synapse. We used a 3 ms and 30 ms step depolarization which has been demonstrated in the prehearing calyx to deplete the RRP and total releasable pool (Lee et al., 2012; Chen et al., 2015). The 3 ms step depletes all those SVs within 60 nm of Cav2 channels while the 30 ms depletes the entire pool of fusion competent SVs within 150-200 nm of Cav2 (Chen et al., 2015). In response to the 3 and 30 ms step depolarizations at P9-11, we found a decrease in the Ca^2+^ charge in the absence of CAST and ELKS (2.66 ±0.2 pC, n = 7, *Cast^−/−^ Elks ^−/−^* vs 3.64 ± 0.27 pC, n = 10, *Cast^+/+^/Elks^+/+^*; p =0.015, t-test, Fig. 9E). However, we found no change in the 3 ms or 30 ms peak ESPC amplitudes compared to control (Fig. 9D, Table 8) and found no difference in the 10-90% EPSC rise time. (Fig. 9C, Table 8). To determine if there was any change in the kinetics of release we normalized the 3 ms and the 30 ms EPSC waveforms. Analysis of the 3 and 30 ms waveforms revealed no change in release kinetics in the *Cast^−/−^/Elks^−/−^* calyces. Taken together, deletion of CAST/ELKS did not result in changes in Cav2 to SV coupling.

**Figure 9.**
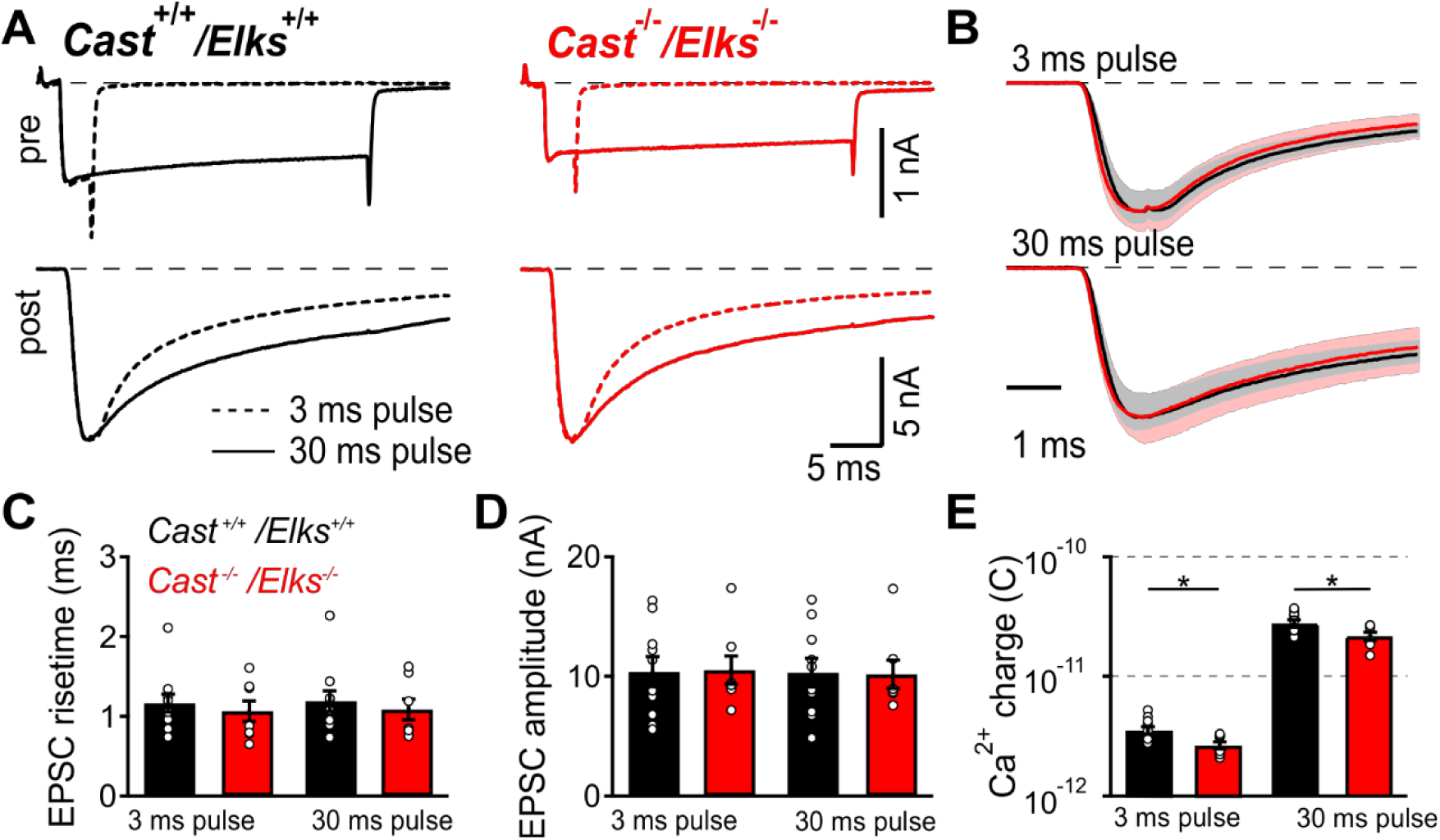
Loss of CAST/ELKS proteins does not change size nor kinetics of release of RRP and total releasable pool. (A) Representative recordings of calcium currents (top) as a response to square pulse depolarization of membrane in the duration of 3 (dotted line) and 30 ms (full line) and EPSC (bottom) recorded as a response to calcium currents in the upper panel. (B) Normalized EPSC traces for 3 ms (top) and 30 ms (bottom). (C, D, E) Mean EPSC amplitude, rise time and ICa charge for *Cast^+/+^ Elks^+/+^* and *Cast^−/−^Elks^−/−^* calyces. Data are represented as mean ± SEM, *Cast^+/+^ Elks^+/+^* n = 10, *Cast^−/−^Elks^−/−^* n = 7, t-test.

**Table 8.** Paired recordings data.

| 3 ms |  |  |  |  |  | 30 ms |  |  |  |
| --- | --- | --- | --- | --- | --- | --- | --- | --- | --- |
| Parameter(unit) | Mean | SEM | (n) | P | Stat. | Mean | SEM | (n) | P |
| **Ca<sup>2+</sup> charge (pC)** |  |  |  |  |  |  |  |  |  |
| *Cast<sup>+/+</sup>/Elks<sup>+/+</sup>* | 3.64 | 0.27 | 10 | control | t-test | 28.11 | 1.9 | 10 | control |
| *Cast<sup>-/-</sup>/Elks<sup>-/-</sup>* | 2.66 | 0.2 | 7 | 0.015 (*) |  | 21.65 | 1.75 | 7 | 0.03(*) |
| **Ca<sup>2+</sup> current amplitude (nA)** |  |  |  |  |  |  |  |  |  |
| *Cast<sup>+/+</sup>/Elks<sup>+/+</sup>* | 1.16 | 0.08 | 10 | control | t-test | 1.16 | 0.07 | 10 | control |
| *Cast<sup>-/-</sup>/Elks<sup>-/-</sup>* | 0.9 | 0.05 | 7 | 0.02 (*) |  | 0.9 | 0.05 | 7 | 0.013 (*) |
| **EPSC amplitude (nA)** |  |  |  |  |  |  |  |  |  |
| *Cast<sup>+/+</sup>/Elks<sup>+/+</sup>* | 10.38 | 1.23 | 10 | control | t-test | 10.3 | 1.2 | 10 | control |
| *Cast<sup>-/-</sup>/Elks<sup>-/-</sup>* | 10.54 | 1.28 | 7 | 0.9 (ns) |  | 10.19 | 1.27 | 7 | 0.94 (ns) |
| **EPSC 10%-90% risetime (ms)** |  |  |  |  |  |  |  |  |  |
| *Cast<sup>+/+</sup>/Elks<sup>+/+</sup>* | 1.17 | 0.12 | 10 | control | t-test | 1.19 | 0.13 | 10 | control |
| *Cast<sup>-/-</sup>/Elks<sup>-/-</sup>* | 1.1 | 0.14 | 7 | 0.59(ns) |  | 1.09 | 0.14 | 7 | 0.61 (ns) |

\*EPSC- excitatory postsynaptic current.

## Discussion

By deleting both CAST and ELKS at an immature glutamatergic presynaptic terminal in developing neuronal circuit, we uncovered that CAST/ELKS regulates presynaptic growth, AZ size and presynaptic Cav2 subtype family current. However, we found synaptic transmission was not impaired. Therefore, we propose that early in synapse development CAST/ELKS have important roles in the morphological development of the presynaptic terminal and regulates presynaptic levels of Cav2 subtype family.

### A developmental role for CAST/ELKS in presynaptic growth and AZ

In general, CAST/ELKS regulation of presynaptic development at a mammalian CNS synapse with conventional AZs has been largely unexplored (Torres and Inestrosa, 2018) and our previous study focused solely on the mature calyx (Dong et al., 2018). Furthermore, *in vitro* hippocampal GABAergic and glutamatergic cultured synapses from neurons lacking CAST/ELKS *in vitro* conditions do not mimic the native environment which may obscure synapse development phenotypes and did not analyze the early stages of synapse development (Liu et al., 2014; Held et al., 2016). Since ELKS levels were significantly reduced at the P7 calyx and undetectable at P9, in contrast to published data in other brain areas that reported ELKS half-life in the range of 11-13 days (Fornasiero et al., 2018), our data indicate that at the calyx of Held ELKS has a half-life of few days (∼3-4 days). These discrepancies can be explained by the facts that our measurements were performed with immunohistochemical method that could result in some amount of the signal not being detected, also protein half-lives can differ in the various brain regions (Fornasiero et al., 2018) and different cellular environments (Dorrbaum et al., 2018) i.e, *in vitro* vs *in vivo*.

How does CAST/ELKS regulate presynaptic development? The mammalian CAST and ELKS proteins bind liprin (Ko et al., 2003), Piccolo/Bassoon (Ohtsuka et al., 2002; Deguchi-Tawarada et al., 2004), RIMs (Ohtsuka et al., 2002; Wang et al., 2002), Munc13-1 (Wang et al., 2009) and syntenin (Ko et al., 2006) to form a large macromolecular complex in the AZ. Therefore, there are multiple possibilities by which loss of CAST/ELKS could impact presynaptic development. Precursor transport vesicles are critical for the assembly of the presynaptic active zone (Gundelfinger et al., 2015) and CAST/ELKS have been found as part of one class of precursor transport vesicles that also contain Piccolo and Bassoon (Maas et al., 2012). Since Bassoon/Piccolo play multiple roles in synapse development (Gundelfinger et al., 2015), it is possible that loss of CAST/ELKS impacts Basson/Piccolo dependent pathways controlling AZ development. CAST/ELKS may regulate presynaptic development through a signaling cascade via the CAST/ELKS liprin interaction (Ko *et al.*, 2003; Dai *et al.*, 2006). Prior studies demonstrated that liprin mutations in *Drosophila* larval NMJ synapses (Kaufmann et al., 2002) and *C. elegans en passant* synapses result increased AZ area (Zhen and Jin, 1999) which is a similar phenotype to the increase in AZ area we detected. In addition, CAST/ELKS interacts with syntenin (Ko et al., 2006) a molecule that is implicated in regulation of synaptic adhesion and intracellular signaling (Beekman and Coffer, 2008). Therefore, this interaction may also contribute to presynaptic development. Finally, it is possible that CAST/ELKS role in presynaptic growth related to its regulation of the transcription factor NF-κB. ELKS was identified as a necessary factor for activation of NF-κB signaling in HeLa cells (Ducut Sigala et al., 2004) and NF-κB has been implicated in synaptogenesis (Boersma et al., 2011).

In mammalian ribbon synapses, loss of CAST (tom Dieck et al., 2012), and combined loss of CAST/ELKS (Hagiwara et al., 2018), resulted in shorter AZs length. In addition, loss of CAST/ELKS results in retinal synapse loss (Hagiwara et al., 2018). However, it is important to note that mechanisms regulating mammalian ribbon synapses development and maintenance are distinct from mammalian synapses with conventional AZs (Lagnado and Schmitz, 2015; Wichmann and Moser, 2015). Similar phenotypes were observed in loss of function studies of *bruchpilot* at *Drosophila*, T-bar synapses (Kittel et al., 2006; Wagh et al., 2006; Fouquet et al., 2009). In addition to the uniqueness of the T-bar structure in Drosophila synapses, the *bruchpilot* C terminal region has little sequence homology known to AZ proteins and is essential to rescue *bruchpilot* defects (Kittel et al., 2006; Wagh et al., 2006; Fouquet et al., 2009).

The rodent calyx of Held undergoes a period of rapid growth expansion from P1 to P9 (Kandler and Friauf, 1993; Hoffpauir et al., 2006; Holcomb et al., 2013) and at the onset of hearing (P12), undergoes morphological remodeling from a cup-like structure to a highly fenestrated structure (Kandler and Friauf, 1993; Kil et al., 1995). In parallel, the AZ area increases and then becomes smaller after hearing onset (Taschenberger et al., 2002). Currently, our understanding of the molecular mechanisms that control these developmental pathways are in their early stages (Baydyuk et al., 2016). Based on our data presented here and our previous study at the mature calyx which demonstrated no change in calyx morphology (Dong et al., 2018) this suggests that CAST/ELKS has a developmental stage specific regulatory role. This is in contrast to other genetic studies that manipulated calyx growth early in development by either ablating BMP1(Xiao et al., 2013) or dynamin (Fan et al., 2016) and led to calyx morphology defects throughout all developmental stages. Also, CAST/ELKS deletion could alter globular bushy cell biology which impacts calyx of Held development of through unknown pathways. Regardless, this suggests that pathways regulated by CAST/ELKS that control presynaptic and AZ growth maybe suppressed by pathways that control calyx remodeling after the onset of hearing.

### CAST/ELKS control Cav2 subtype channels levels at the presynaptic AZ

Since we found a reduction in all Cav2 currents in the absence of CAST/ELKS this indicates the mechanisms by which CAST/ELKS regulates presynaptic Cav2 levels is similar between the different Cav2 subtypes. Furthermore it indicates that CAST/ELKS regulation is not specific to regulation Cav2.1 channels or only at the mature calyx of Held (Dong *et al.*, 2018). Using SDS-FFRIL our reduction in Cav2.1 currents was found to be correlated with a reduction in Cav2.1 numbers, however we were unable correlate the reduction in Cav2.2 and 2.3 currents with SDS-FFRIL. Unfortunately, current available antibodies against Cav2.2 α_1_ subunit are not specific for the Cav2.2 for use with SDS-FFRIL (data not shown) and Cav2.3 antibodies suitable for SDS-FFRIL are not available. Therefore, since the reduction in Cav2.2 and 2.3 currents levels was affected in a similar manner we assumed that Cav2.2 and Cav2.3 numbers were also reduced. Development of Cav2.2 and Cav2.3 α_1_ subunit antibodies compatible with SDS-FFRIL will be needed for future experiments counting Cav2.2 and Cav2.3 channel numbers.

CAST/ELKS has been reported to directly interact with the linker II-III domain of the Cav2.1 α_1_ subunit and with the Cavβ4 (Kiyonaka et al., 2012). Since the domain II-III linker region has 65% consensus between the Cav2 α_1_ subunits (Zamponi, 2016) we favor a model by which CAST/ELKS regulates presynaptic Cav2 levels through Cavβ interactions or other binding partners. Previous reports have demonstrated that in retinal ribbon synapses loss of CAST results in reduction of Cav1.4 channels while loss of CAST/ELKSs resulted in a reduction in Cav1.4 current density (Hagiwara et al., 2018). Furthermore, the conservation between Cav2 and Cav1.4 domain II-III linkers region is poor (Zamponi, 2016). CAST/ELKS interacts with RIM1/2 PDZ with extremely high affinity (Lu et al., 2005) and Cav2 current density is selectively reduced with RIMs deletion (Han et al., 2011; Kaeser et al., 2011). Therefore, it is possible that RIM and CAST/ELKS interaction with Cavβ forms a macromolecular complex that controls Cav2 abundance.

### Loss of CAST/ELKS does not impact synaptic transmission

Despite leading to dramatic changes in presynaptic morphology and a reduction in Cav2 currents and numbers, CAST/ELKS loss did not impact AP-evoked synaptic transmission. Although glutamate receptor desensitization seen at the immature calyx (Schneggenburger et al., 1999; Renden et al., 2005) can impact accuracy of RRP measurements, we used 1mM Kyn which previously has been shown to enable an accurate RRP estimates (Lee et al., 2008). Furthermore our paired recordings were done in the presence of 2mM Kyn and CTZ to block all desensitization and we found no difference in the RRP size as measured by our 3 ms step depolarization. In addition, our paired recording data rules out that changes in Cav2 channel activation play a role and that coupling is impacted. This is in contrast to clear defects at the mature calyx which reported a reduction in the RRP size, increase in *Pr*, (Dong et al., 2018) and defects seen at cultured hippocampal neurons (Liu et al., 2014; Held et al., 2016). Therefore, this indicates that at the immature calyx the loss of CAST/ELKS can be compensated regarding synaptic transmission. Therefore, it is possible that the morphological defects mask the roles of CAST/ELKS in regulating synaptic transmission. In this scenario, the reduction in calyx volume may result in similar or increased intra-terminal calcium concentration levels compared to wild-type terminals to counter the reduction in calcium channels. In addition, the increase in the number of docked vesicles may also contribute to the lack of synaptic transmission phenotype.

In summary, our data have identified a role for CAST/ELKS in presynaptic maturation of a developing neuronal circuit. Exact mechanisms of action of CAST/ELKS in regulating abundance of Cav2 levels and morphological development of calyx of Held and their role in other CNS synapses needs to be further investigated as well as their involvement in neurological disorders caused by alterations in neurodevelopment.

## Material and methods

### Ethical approval

All animals were used in accordance with animal welfare laws approved by the Institutional Committee for Care and Use of Animals at the University of Yamanashi, the Max Planck Florida Institute for Neuroscience, and the University of Iowa. Animals were housed with 12 h light/dark cycle and *ab libitum* food/water supply. Animals of both sexes were used for all experiments. Age of animals was ranging from P7 to P11. *Cast^−/−^ Elks^fl/fl^* mice were generated as previously described (Dong et al., 2018). All available measures were taken to minimize animals pain and suffering.

## EXPERIMENTAL DESIGN

### Recombinant Viral Vector Production

Helper Dependent Adenoviral (HdAd) vectors expressing Cre recombinase were produced as previously described (Montesinos et al., 2016; Dong et al., 2018). In brief, expression cassette with Cre recombinase was cloned into the pdelta28E4 plasmid using *Asc*I enzyme digestion site. Plasmid has been modified to also contain a separate EGFP or myristolated EGFP (mEGFP) expression cassette. All expression cassettes were driven by the 470 bp hsyn promoter and the final HdAd plasmid allows for expression of Cre independently of EGFP. Then, the pHAD plasmid was linearized by *PmeI* enzyme to expose the ends of the 5’ and 3’ inverted terminal repeats (ITRs) and transfected into 116 producer cells (Profection® Mammalian Transfection System, Promega, Madison, WI, USA). Helper virus (HV) was added the following day for HdAd production. Forty-eight hours post infection, after cytopathic effects have taken place, cells were subjected to three freeze/thaw cycles for lysis and release of the viral particles. HdAd was purified by CsCl ultracentrifugation. HdAd was stored at −80 °C in storage buffer (10 mM HEPES, 1 mM MgCl_2_, 250 mM sucrose, pH 7.4).

### Stereotaxic injection

Stereotactic injection of HdAds into the cochlear nucleus (CN) was performed at postnatal day 1 (P1), as described (Chen et al., 2013). Briefly, animals were anesthetized by hypothermia for 5 mins and then injected with 1.5 μL HdAd expressing codon-optimized Cre recombinase and EGFP (1×10^7^ transducing units/l). After surgery, animals were allowed to recover under a heating lamp before being returned to their respective cages.

### Electrophysiological recordings

#### Slice preparation

Acute brain slices were prepared as previously described (Chen et al., 2013). Briefly, after decapitation of the animal, the brain was immersed in low-calcium artificial cerebrospinal fluid (aCSF) containing (in mM): 125 NaCl, 2.5 KCl, 10 glucose, 25 NaHCO_3_, 1.25 NaH_2_PO_4_, 0.4 L-ascorbic acid, 3 myo-inositol, 2 Na-pyruvate, 3 MgCl_2_ and 0.1 CaCl_2_ continuously bubbled with 95% O_2_-5% CO_2_ (pH 7.3). Acute coronal brainstem slices (200 μm) containing the MNTB were obtained using a Leica VT 1200S vibratome. Slices were immediately transferred to an incubation beaker containing normal aCSF (same as the low-calcium aCSF except that 1 mM MgCl_2_ and 1 or 2 mM CaCl_2_ were used) at 37°C, continuously bubbled with 95% O_2_-5% CO_2_. The slices were allowed for recovery for 45-60 min and then were transferred to a recording chamber with the same aCSF solution at room temperature (≈25°C).

#### Presynaptic Ca^2+^ current recordings

Presynaptic Ca^2+^ currents were isolated by applying 1 µM tetrodotoxin (TTX; Tocris Bioscience), 100 µM 4-aminopyridine (4-AP; Tocris) and 20 mM tetraethylammonium chloride (TEA-Cl; Sigma Aldrich) with aCSF to block Na^+^ and K^+^ channels. Current-voltage (IV) relationships were recorded in the presence 1 mM CaCl_2_, while pharmacological isolation of VGCC subtypes was performed in 2 mM CaCl_2_. To measure VGCC subtypes, 200 nM ω-agatoxin IVA (Alomone labs) that blocks Cav2.1, 2 µM ω-conotoxin GVIA (Alomone labs) that blocks Cav2.2 and 50 µM CdCl_2_ to block the remaining I_Ca_ were applied. All subtype specific toxins were supplemented with 0.1 mg/mL cytochrome C. Presynaptic patch pipettes were filled with (mM): 145 Cs-gluconate, 20 TEA-Cl, 10 HEPES, 5 Na_2_-phosphocreatine, 4 MgATP, 0.3 NaGTP and 0.5 EGTA. Calyces were held at −80 mV and 10 ms step depolarization from −80 mV to 0 mV were applied. Rs was typically <20 MΩ and online compensated to 6 MΩ. Leak current was < 100 pA.

#### Paired pre- and postsynaptic recordings

External Ca^2+^ concentration was 2 mM for paired recordings, 2 mM kynurenic acid, 100 μM cyclothiazide (Tocris Bioscience), 50 μM D-AP5, 100 μM 4-AP, 20 mM TEA and 1 μM TTX were included in external solution. Presynaptic calyx terminals and postsynaptic principal neurons of medial nucleus of trapezoid body (MNTB) were simultaneously whole-cell voltage-clamped at −80 mV and −60 mV, respectively. Patch pipettes had an open tip resistance of 5-6 MΩ and 3-4 MΩ for presynaptic and postsynaptic recordings, respectively. Presynaptic R_S_ (<15 MΩ) was online compensated to 8 MΩ. Postsynaptic R_S_ (<8 MΩ) was online compensated to 3 MΩ, and remaining R_S_ was compensated offline to 0.

#### Afferent fiber stimulation

Afferent fiber stimulation was performed as described previously (Forsythe and Barnes-Davies, 1993a, b). Briefly, a bipolar electrode was placed between brainstem midline and MNTB. MNTB principal neurons were whole-cell voltage-clamped at −60 mV. 2 mM external Ca^2+^ were used for recording on P9-11 calyces. Evoked EPSCs were recorded with the use of 1 mM kynurenic acid (Tocris Bioscience) to minimize receptor saturation (Trussell, 1998); 50 μM D-AP-5 (Tocris Bioscience) to block NMDA receptor, 20 μM bicuculline (Tocris Bioscience) and 5 μM strychnine (Tocris Bioscience) to block inhibitory postsynaptic currents (IPSCs). Patch pipettes had open tip resistances of 3-4 MΩ and were filled with (in mM): 145 Cs-gluconate, 20 TEA-Cl, 10 HEPES, 5 Na_2_-phosphocreatine, 4 MgATP, 0.3 NaGTP, 6 QX-314, and 5 EGTA. Series resistance (R_S_, 3-8 MΩ) were online compensated to 3 MΩ and the remaining R_S_ was offline compensated to 0 MΩ for all EPSCs (Traynelis, 1998).

#### Miniature postsynaptic currents (mEPSCs)

External Ca^2+^ of 2 mM was used for mEPSCs recording. MNTB principal neurons were whole-cell voltage-clamped at −80 mV. ACSF was supplemented with 50 μM D-AP5, 20 μM bicuculline, 5 μM strychnine, 1 μM TTX and 20 mM TEA. Rs (<8 MΩ) was not compensated.

#### Immunohistochemistry

Mice were anesthetized with an intraperitoneal (i.p.) injection of tribromoethanol (250 mg/kg body weight) and transcardially perfused with ice-cold 0.1 M phosphate buffer (PB) (pH 7.4). Brains were removed and rapidly frozen in optimal cutting temperature compound (OCT) using dry ice and 100% ethanol. 8-µm-thick coronal sections of the brainstem were obtained using a cryostat HM620, thaw mounted on microscope slides (Fisherbrand^TM^ Superfros^TM^ Plus, Fisher Scientific) and air dried for 15 min. Sections were fixed by immersion in ice-cold 95% ethanol at 4°C for 30 min followed by acetone for 1 min at RT. Prior to the staining, sections were washed in 0.1 M PB and incubated 20 min at RT in 0.1 M PB blocking solution containing 2% normal goat serum (NGS) and 0.2% Triton X-100. Afterwards, sections were incubated at 4°C overnight with the primary antibodies diluted in the same blocking solution (1:100 (1 mg/ml), Rabbit anti-CAST, and 1:100 (1mg/ml) Rabbit anti-ELKS, antibodies made in the lab of Dr. Ohtsuka, T99; 1:100, polyclonal Guinea pig anti-VGLUT1, Synaptic Systems, RRID: AB_887878).

Subsequently, sections were rinsed 3×10 min in 0.1 M PB and incubated with the secondary antibodies diluted in 0.1 M PB containing 0.1% Triton X-100 for 2 h at RT 1:200 (Cy^TM^ 2 AffiniPure goat anti-rabbit IgG (H+L), Jackson Immunoresearch, RRID: AB_2338021; 1:200, Alexa Fluor® 647-conjugated AffiniPure donkey anti-guinea pig, Jackson Immunoresearch, RRID: AB_2340476). After 3×10 min in 0.1 M PB washed sections were mounted using Fluoromount-G® mounting medium (SouthernBiotech).

#### Presynaptic terminal reconstructions

Stereotactic injection of mice at P1 was performed using Cre mEGFP expressing virus and at P9 mice were anesthetized with an intraperitoneal (i.p.) injection of tribromoethanol (250 mg/kg body weight) and transcardially perfused with ice-cold 0.1 M phosphate buffer (PB) (pH 7.4) followed by perfusion with 4% paraformaldehyde (PFA). Brains were removed and left overnight in the same PFA solution. Next day brains were sliced on Leica VT1200 vibratome into 40 µm thin sections. mEGFP positive slices were identified and mounted on cover slips using Aqua Polymount solution (Polysciences, Inc.).

### Electron microscopy

#### Preembedding immuno-electron microscopy

Preembedding immuno-electron microscopy procedure was described in detail in (Montesinos et al., 2015; Dong et al., 2018). Briefly, wild type (C57/BL6J) and Cre virus injected *Cast*^−/−^/*Elks*^fl/fl^ P9 mice were anesthetized and perfused transcardially with phosphate-buffered saline (PBS, 150 mM NaCl, 25 mM Sørensen’s phosphate buffer, pH 7.4) followed by fixative solution for 7-9 min containing 4% PFA, 0.5% glutaraldehyde, and 0.2% picric acid in 100 mM Sørensen’s phosphate buffer (PB, pH 7.4). Brains were post-fixed with 4% PFA in PB overnight and 50-µm coronal sections of the brainstem were obtained on a vibratome (Leica VT1200S). Expression of EGFP at calyx of Held terminals was visualized using an epifluorescence inverted microscope (CKX41, Olympus) equipped with an XCite Series 120Q lamp (Excelitas technologies), and only those samples showing EGFP were further processed. After washing with PB, sections were cryoprotected with 10%, 20% and 30% sucrose in PB and submersed into liquid nitrogen, then thawed. Sections were incubated in a blocking solution containing 10% normal goat serum (NGS), 1% fish skin gelatin (FSG), in 50 mM Tris-buffered saline (TBS, 150 mM NaCl, 50 mM Tris, pH 7.4) for 1h, and incubated with an anti-GFP antibody (0.1µg/ml, ab6556, Abcam, RRID: AB_305564) diluted in TBS containing 1% NGS, 0.1% FSG plus 0.05% NaN_3_ at 4°C for 48h. After washing with TBS, sections were incubated overnight in nanogold conjugated goat anti-rabbit IgG (1:100, Nanoprobes, RRID: AB_2340585) diluted in TBS containing 1% NGS and 0.1% FSG. Immunogold-labeled sections were washed in PBS, briefly fixed with 1% glutaraldehyde in PBS, and silver intensified using HQ silver intensification kit (Nanoprobe). After washing with PB, sections were treated with 0.5% OsO_4_ in 0.1M PB for 20 min, en-bloc stained with 1% uranyl acetate, dehydrated and flat embedded in Durcupan resin (Sigma-Aldrich). After trimming out the MNTB region, ultrathin sections were prepared with 40 nm-thickness using an ultramicrotome (EM UC7, Leica). Sections were counterstained with uranyl acetate and lead citrate and examined in a Tecnai G2 Spirit BioTwin transmission electron microscope (Thermo Fisher Scientific) at 100kV acceleration voltage. Images were taken with a Veleta CCD camera (Olympus) operated by TIA software (Thermo Fisher Scientific). Images used for quantification were taken at 60,000x magnification.

#### SDS-digested freeze fracture replica immuno labeling (SDS-FFRIL)

Mice were injected with Cre virus harboring myristoylated EGFP at P1. P9 animals were anesthetized with tribromoethanol (250 mg/kg of body weight, i.p.) and perfused transcardially with PBS followed by 2% PFA, and 0.2% picric acid in 0.1 M PB for 12 min. 130-µm-thick coronal sections of the brainstem were obtained using a vibratome (Leica VT1200S). Only those samples showing expression of myristoylated EGFP were selected. The brain slices containing the MNTB region were cryoprotected in 10%, 20% and 30% glycerol at 4°C overnight. Small pieces containing MNTB region were trimmed out and frozen using a high-pressure freezing machine (HPM100, Leica). Frozen samples were fractured in two halves using a double replication device at −120°C, replicated first with a 2 nm carbon deposition, shadowed from 60-degree angle by carbon-platinum of 2 nm and supported by a final carbon deposition of 20-30nm, using a JFDV Freeze fracture machine (JEOL/Boeckeler). The tissue was dissolved by placing the replicas in a digesting solution containing 2.5% SDS, 20% sucrose and 15 mM Tris-HCl (pH 8.3) with gentle agitation in an oven at 82.5°C for 18 hours. The replicas were washed and blocked with 4% BSA and 1% FSG in TBS for 1 hour, then incubated with a mixture of primary antibody; rabbit anti-GFP (1 µg/ml, AbCam, cat# ab6556, RRID: AB_3055564) and guinea pig anti-α1 subunit of Cav2.1 (0.7 µg/ml, Synaptic Systems, cat# 152205, RRID:AB_2619842) diluted in 0.04% BSA and 0.01% FSG in TBS for 18 h at room temperature. After several washes, the replicas were incubated in donkey anti-rabbit IgG conjugated to 6 nm gold particles and donkey anti-guinea pig IgG conjugated to 12 nm gold particles (1:30 respectively; Jackson Immunoresearch, RRID: AB_2340609 RRID: AB_2340465) diluted in 0.04% BSA and 0.01% FSG in TBS for 18 h at room temperature. After being washed, the replicas were picked up on 100-parallel-bar copper grids or aperture grids and examined with a Tecnai G2 Spirit BioTwin transmission electron microscope (Thermo Fisher Scientific) at 100 kV acceleration voltage. Images were taken with a Veleta CCD camera (Olympus).

## QUANTIFICATION AND STATISTICAL ANALYSIS

### Confocal imaging and image analysis

Confocal images were acquired with a Zeiss LSM 780 and Zeiss LSM 700 confocal scanning microscope. Confocal scans for each fluorochrome was acquired sequentially using a oil immersed 63x/1.3(NA) apochromat MultiImmersion Zeiss objective. The intensity emission signal from each channel was adjusted to below saturation level. Stack images were collected using 0.44 microns plane line scans with line average of 4 times. Images were processed with Fiji imaging analysis software (http://fiji.sc., RRID:SCR_002285). For signal intensity calculations, ROI of a single plane image was detected by using Wand tool that creates a selection by tracing objects/pixels of uniform signal intensities. Visual inspection was used to determine that selection was precise and constrained to Vgut1 signal. Same ROI was added to image with ELKS signal and using Measurement option in Analysis mean signal intensities were obtained and divided (ELKS/VGLUT1) for ratio calculations. Calyx reconstructions were done using Imaris Measurement Pro (BitPlane) by combination of manual and automatic signal detection of single planes from Z-stack confocal images.

### TEM image analysis

We identified calyces which were positive for Cre expression by immunogold labeling with an anti-GFP antibody and compared EGFP-positive terminals (*Cast^−/−^/Elks^−/−^)* to EGFP-negative terminals (*Cast^−/−^*) in the same slice or to calyces in the wild type sample. All TEM data were analyzed using Fiji imaging analysis software (Schindelin et al., 2012). Each presynaptic active zone (AZ) was defined as the membrane directly opposing postsynaptic density, and the length of each AZ was measured. Vesicles within 200 nm from each AZ were manually selected and their distances relative to the AZ were calculated using a 32-bit Euclidean distance map generated from the AZ. For data analysis, vesicle distances were binned every 5 nm and counted (Montesinos et al., 2015; Dong et al., 2018). Vesicles less than 5 nm from the AZ were considered “docked”(Taschenberger et al., 2002). Three animals for each condition and 40 AZs per animal were analyzed.

### SDS-FFRIL image collection and analysis

Presynaptic P-faces (protoplasmic face of the cell membrane) of the calyx of Held were morphologically identified by existence of cross-fractured cytoplasm containing synaptic vesicles and/or by existence of glutamate receptor clusters uniquely shown on the adjacent E-face (exoplasmic face) of the postsynaptic cell of principle neuron of MNTB (Budisantoso et al., 2012; Nakamura et al., 2015). Specifically, immunogold labeled presynaptic P-faces that were in contact with the postsynaptic soma were imaged at 43,000x magnification. EGFP positive presynaptic P-faces were identified by the existence of 6-nm immunogold particles as *Cast^−/−^/Elks^−/−^*, and EGFP negative calyx terminals were categorized as *Cast^−/−^*. We analyzed the distribution and number of Cav2.1 from the 12 nm immunogold labeling in both EGFP positive and negative terminals. Three P-faces from the replicas of *Cast^−/−^* and *Cast^−/−^/Elks^−/−^* samples (6 total), including small to large pieces, were analyzed comprising 45.8 and 56.6 µm^2^ of P-face area, respectively. For each continuous P-face, images were manually stitched and minimally adjusted for brightness and contrast (Adobe Photoshop CS6). 12-nm gold particles corresponding to Cav2.1 channels were thresholded and quantified using Microscopy Image Browser (MIB) (Belevich et al., 2016). A custom macro was used in Fiji to draw a circle with a 30- and 100-nm radius from each 12-nm gold particle. Overlapping 30-nm circles were considered clusters of closely distributed Cav2.1 channels (Nakamura et al., 2015). Clusters and the particles within them were then analyzed.

### Electrophysiological data analysis

Electrophysiological data were analyzed offline with Fitmaster (Heka, RRID:SCR_016233) and custom functions written in Igor Pro (version 6.36; Wavemetrics, RRID:SCR_000325, Patchers Power Tools, RRID:SCR_001950).

Peak Ca^2+^ current-voltage relationships were fitted according to a Hodgkin-Huxley formalism assuming four independent gates and Goldman-Hodgkin-Katz for open-channel conductance as a function of voltage Γ(V):

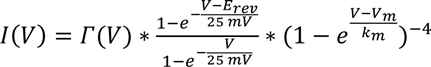

with E_rev_ as reversal potential, V_m_ as half-maximal activation voltage per gate, and k_m_ as the voltage-dependence of activation. Tail currents were measured as peaks minus baseline, plotted as a function of voltage and fitted with a Boltzmann function:

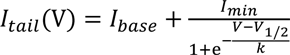

where V_1/2_ represents the half-maximal voltage and k the corresponding slope factor.

EPSC amplitudes were measured as peak minus baseline. The size of the readily-releasable pool (RRP) was determined from 100 Hz train stimulation using the back-extrapolation method as described previously (Schneggenburger *et al.*, 1999; Neher, 2015).

For analysis of mEPSCs custom written script by H. Taschenberger (Clements & Bekkers, 1997) in Igor Pro was used.

## STATISTICS

Statistics were performed with Prism 8 (GraphPad Software, RRID:SCR_002798). Shapiro-Wilk test was used to test for the assumption of normal distribution and Bartlett’s test to test for assumption of equal variance among groups. To compare two normally distributed groups, an unpaired two-tailed Student’s t test was used. To compare more than two normally distributed groups, a one-way ANOVA with a post hoc Dunnett’s test was used. To compare not normally distributed groups a Kruskal-Wallis with a post hoc Dunn’s test was used. Data are reported as mean ± SEM or cumulative distribution. Statistical significance was accepted at p < 0.05.

## Acknowledgements

We thank Rachel Satterfield for HdAd viral vector production, Dr. Stacia Phillips for editing, We thank Dr.Henrique von Gersdorff for comments on the manuscript. We thank Dr. Phillip Ng and Dr. Brendan Lee for gifts of HdAd packing plasmids and HdAd stuffer DNA, respectively and Dr. Holger Taschenberger for sharing custom mEPSC analysis routine.

## Author contributions

W.D., T.R., R.O.G., C.I.T., P.V., M.S.M., D.G.G, K.G., M.L., N.K., carried out experiments and analyzed data. T.O. developed mouse model. T.O. and S.M.Y, Jr planned the project and analyzed data. S.M.Y, Jr. wrote the manuscript. All authors jointly revised the paper.

## References

Ackermann F, Waites CL, Garner CC (2015) Presynaptic active zones in invertebrates and vertebrates. EMBO Rep 16:923–938.

Althof D, Baehrens D, Watanabe M, Suzuki N, Fakler B, Kulik A (2015) Inhibitory and excitatory axon terminals share a common nano-architecture of their Cav2.1 (P/Q-type) Ca(2+) channels. Frontiers in cellular neuroscience 9:315.

Baydyuk M, Xu J, Wu LG (2016) The calyx of Held in the auditory system: Structure, function, and development. Hear Res 338:22–31.

Beekman JM, Coffer PJ (2008) The ins and outs of syntenin, a multifunctional intracellular adaptor protein. Journal of cell science 121:1349–1355.

Belevich I, Joensuu M, Kumar D, Vihinen H, Jokitalo E (2016) Microscopy Image Browser: A Platform for Segmentation and Analysis of Multidimensional Datasets. PLoS Biol 14:e1002340.

Boersma MC, Dresselhaus EC, De Biase LM, Mihalas AB, Bergles DE, Meffert MK (2011) A requirement for nuclear factor-kappaB in developmental and plasticity-associated synaptogenesis. J Neurosci 31:5414–5425.

Borst JG, Sakmann B (1996) Calcium influx and transmitter release in a fast CNS synapse. Nature 383:431–434.

Budisantoso T, Matsui K, Kamasawa N, Fukazawa Y, Shigemoto R (2012) Mechanisms underlying signal filtering at a multisynapse contact. J Neurosci 32:2357–2376.

Chen Z, Cooper B, Kalla S, Varoqueaux F, Young SM, Jr. (2013) The Munc13 proteins differentially regulate readily releasable pool dynamics and calcium-dependent recovery at a central synapse. The Journal of neuroscience : the official journal of the Society for Neuroscience 33:8336–8351.

Chen Z, Das B, Nakamura Y, DiGregorio DA, Young SM, Jr. (2015) Ca2+ channel to synaptic vesicle distance accounts for the readily releasable pool kinetics at a functionally mature auditory synapse. J Neurosci 35:2083–2100.

Cortes-Saladelafont E, Lipstein N, Garcia-Cazorla A (2018) Presynaptic disorders: a clinical and pathophysiological approach focused on the synaptic vesicle. J Inherit Metab Dis 41:1131–1145.

Deguchi-Tawarada M, Inoue E, Takao-Rikitsu E, Inoue M, Ohtsuka T, Takai Y (2004) CAST2: identification and characterization of a protein structurally related to the presynaptic cytomatrix protein CAST. Genes to cells : devoted to molecular & cellular mechanisms 9:15–23.

Deken SL, Vincent R, Hadwiger G, Liu Q, Wang ZW, Nonet ML (2005) Redundant localization mechanisms of RIM and ELKS in Caenorhabditis elegans. J Neurosci 25:5975–5983.

Di Guilmi MN, Wang T, Inchauspe CG, Forsythe ID, Ferrari MD, van den Maagdenberg AM, Borst JG, Uchitel OD (2014) Synaptic gain-of-function effects of mutant Cav2.1 channels in a mouse model of familial hemiplegic migraine are due to increased basal [Ca2+]i. J Neurosci 34:7047–7058.

Dong W, Radulovic T, Goral RO, Thomas C, Suarez Montesinos M, Guerrero-Given D, Hagiwara A, Putzke T, Hida Y, Abe M, Sakimura K, Kamasawa N, Ohtsuka T, Young SM, Jr. (2018) CAST/ELKS Proteins Control Voltage-Gated Ca(2+) Channel Density and Synaptic Release Probability at a Mammalian Central Synapse. Cell reports 24:284–293 e286.

Dorrbaum AR, Kochen L, Langer JD, Schuman EM (2018) Local and global influences on protein turnover in neurons and glia. eLife 7.

Doughty JM, Barnes-Davies M, Rusznak Z, Harasztosi C, Forsythe ID (1998) Contrasting Ca2+ channel subtypes at cell bodies and synaptic terminals of rat anterioventral cochlear bushy neurones. The Journal of physiology 512 (Pt 2):365–376.

Ducut Sigala JL, Bottero V, Young DB, Shevchenko A, Mercurio F, Verma IM (2004) Activation of transcription factor NF-kappaB requires ELKS, an IkappaB kinase regulatory subunit. Science 304:1963–1967.

Fan F, Funk L, Lou X (2016) Dynamin 1- and 3-Mediated Endocytosis Is Essential for the Development of a Large Central Synapse In Vivo. J Neurosci 36:6097–6115.

Fedchyshyn MJ, Wang LY (2005) Developmental transformation of the release modality at the calyx of Held synapse. The Journal of neuroscience : the official journal of the Society for Neuroscience 25:4131–4140.

Fornasiero EF et al. (2018) Precisely measured protein lifetimes in the mouse brain reveal differences across tissues and subcellular fractions. Nat Commun 9:4230.

Forsythe ID, Barnes-Davies M (1993a) The binaural auditory pathway: membrane currents limiting multiple action potential generation in the rat medial nucleus of the trapezoid body. Proceedings Biological sciences / The Royal Society 251:143–150.

Forsythe ID, Barnes-Davies M (1993b) The binaural auditory pathway: excitatory amino acid receptors mediate dual timecourse excitatory postsynaptic currents in the rat medial nucleus of the trapezoid body. Proceedings Biological sciences / The Royal Society 251:151–157.

Fouquet W, Owald D, Wichmann C, Mertel S, Depner H, Dyba M, Hallermann S, Kittel RJ, Eimer S, Sigrist SJ (2009) Maturation of active zone assembly by Drosophila Bruchpilot. The Journal of cell biology 186:129–145.

Gundelfinger ED, Reissner C, Garner CC (2015) Role of Bassoon and Piccolo in Assembly and Molecular Organization of the Active Zone. Frontiers in synaptic neuroscience 7:19.

Hagiwara A, Fukazawa Y, Deguchi-Tawarada M, Ohtsuka T, Shigemoto R (2005) Differential distribution of release-related proteins in the hippocampal CA3 area as revealed by freeze-fracture replica labeling. The Journal of comparative neurology 489:195–216.

Hagiwara A, Kitahara Y, Grabner CP, Vogl C, Abe M, Kitta R, Ohta K, Nakamura K, Sakimura K, Moser T, Nishi A, Ohtsuka T (2018) Cytomatrix proteins CAST and ELKS regulate retinal photoreceptor development and maintenance. The Journal of cell biology 217:3993–4006.

Han Y, Kaeser PS, Sudhof TC, Schneggenburger R (2011) RIM determines Ca(2)+ channel density and vesicle docking at the presynaptic active zone. Neuron 69:304–316.

Held RG, Liu C, Kaeser PS (2016) ELKS controls the pool of readily releasable vesicles at excitatory synapses through its N-terminal coiled-coil domains. eLife 5.

Hoffpauir BK, Grimes JL, Mathers PH, Spirou GA (2006) Synaptogenesis of the calyx of Held: rapid onset of function and one-to-one morphological innervation. J Neurosci 26:5511–5523.

Holcomb PS, Hoffpauir BK, Hoyson MC, Jackson DR, Deerinck TJ, Marrs GS, Dehoff M, Wu J, Ellisman MH, Spirou GA (2013) Synaptic inputs compete during rapid formation of the calyx of Held: a new model system for neural development. J Neurosci 33:12954–12969.

Holderith N, Lorincz A, Katona G, Rozsa B, Kulik A, Watanabe M, Nusser Z (2012) Release probability of hippocampal glutamatergic terminals scales with the size of the active zone. Nat Neurosci 15:988–997.

Iwasaki S, Takahashi T (1998) Developmental changes in calcium channel types mediating synaptic transmission in rat auditory brainstem. The Journal of physiology 509 (Pt 2):419–423.

Kaeser PS, Deng L, Wang Y, Dulubova I, Liu X, Rizo J, Sudhof TC (2011) RIM proteins tether Ca2+ channels to presynaptic active zones via a direct PDZ-domain interaction. Cell 144:282–295.

Kandler K, Friauf E (1993) Pre- and postnatal development of efferent connections of the cochlear nucleus in the rat. The Journal of comparative neurology 328:161–184.

Kaufmann N, DeProto J, Ranjan R, Wan H, Van Vactor D (2002) Drosophila liprin-alpha and the receptor phosphatase Dlar control synapse morphogenesis. Neuron 34:27–38.

Kavalali ET (2014) The mechanisms and functions of spontaneous neurotransmitter release. Nat Rev Neurosci 16:5–16.

Kil J, Kageyama GH, Semple MN, Kitzes LM (1995) Development of ventral cochlear nucleus projections to the superior olivary complex in gerbil. The Journal of comparative neurology 353:317–340.

Kittel RJ, Wichmann C, Rasse TM, Fouquet W, Schmidt M, Schmid A, Wagh DA, Pawlu C, Kellner RR, Willig KI, Hell SW, Buchner E, Heckmann M, Sigrist SJ (2006) Bruchpilot promotes active zone assembly, Ca2+ channel clustering, and vesicle release. Science 312:1051–1054.

Kiyonaka S, Nakajima H, Takada Y, Hida Y, Yoshioka T, Hagiwara A, Kitajima I, Mori Y, Ohtsuka T (2012) Physical and functional interaction of the active zone protein CAST/ERC2 and the beta-subunit of the voltage-dependent Ca(2+) channel. Journal of biochemistry 152:149–159.

Ko J, Na M, Kim S, Lee JR, Kim E (2003) Interaction of the ERC family of RIM-binding proteins with the liprin-alpha family of multidomain proteins. The Journal of biological chemistry 278:42377–42385.

Ko J, Yoon C, Piccoli G, Chung HS, Kim K, Lee JR, Lee HW, Kim H, Sala C, Kim E (2006) Organization of the presynaptic active zone by ERC2/CAST1-dependent clustering of the tandem PDZ protein syntenin-1. J Neurosci 26:963–970.

Lagnado L, Schmitz F (2015) Ribbon Synapses and Visual Processing in the Retina. Annu Rev Vis Sci 1:235–262.

Lee JS, Ho WK, Lee SH (2012) Actin-dependent rapid recruitment of reluctant synaptic vesicles into a fast-releasing vesicle pool. Proceedings of the National Academy of Sciences of the United States of America 109:E765–774.

Lee JS, Kim MH, Ho WK, Lee SH (2008) Presynaptic release probability and readily releasable pool size are regulated by two independent mechanisms during posttetanic potentiation at the calyx of Held synapse. J Neurosci 28:7945–7953.

Lenkey N, Kirizs T, Holderith N, Mate Z, Szabo G, Vizi ES, Hajos N, Nusser Z (2015) Tonic endocannabinoid-mediated modulation of GABA release is independent of the CB1 content of axon terminals. Nat Commun 6:6557.

Liu C, Bickford LS, Held RG, Nyitrai H, Sudhof TC, Kaeser PS (2014) The active zone protein family ELKS supports Ca2+ influx at nerve terminals of inhibitory hippocampal neurons. J Neurosci 34:12289–12303.

Lu J, Li H, Wang Y, Sudhof TC, Rizo J (2005) Solution structure of the RIM1alpha PDZ domain in complex with an ELKS1b C-terminal peptide. Journal of molecular biology 352:455–466.

Maas C, Torres VI, Altrock WD, Leal-Ortiz S, Wagh D, Terry-Lorenzo RT, Fejtova A, Gundelfinger ED, Ziv NE, Garner CC (2012) Formation of Golgi-derived active zone precursor vesicles. J Neurosci 32:11095–11108.

Maschi D, Klyachko VA (2017) Spatiotemporal Regulation of Synaptic Vesicle Fusion Sites in Central Synapses. Neuron 94:65–73 e63.

Miki T, Kaufmann WA, Malagon G, Gomez L, Tabuchi K, Watanabe M, Shigemoto R, Marty A (2017) Numbers of presynaptic Ca(2+) channel clusters match those of functionally defined vesicular docking sites in single central synapses. Proc Natl Acad Sci U S A 114:E5246–E5255.

Mochida S, Hida Y, Tanifuji S, Hagiwara A, Hamada S, Abe M, Ma H, Yasumura M, Kitajima I, Sakimura K, Ohtsuka T (2016) SAD-B Phosphorylation of CAST Controls Active Zone Vesicle Recycling for Synaptic Depression. Cell reports 16:2901–2913.

Montesinos MS, Satterfield R, Young SM, Jr. (2016) Helper-Dependent Adenoviral Vectors and Their Use for Neuroscience Applications. Methods Mol Biol 1474:73–90.

Montesinos MS, Dong W, Goff K, Das B, Guerrero-Given D, Schmalzigaug R, Premont RT, Satterfield R, Kamasawa N, Young SM, Jr. (2015) Presynaptic Deletion of GIT Proteins Results in Increased Synaptic Strength at a Mammalian Central Synapse. Neuron 88:918–925.

Nakamura Y, Harada H, Kamasawa N, Matsui K, Rothman JS, Shigemoto R, Silver RA, DiGregorio DA, Takahashi T (2015) Nanoscale distribution of presynaptic Ca(2+) channels and its impact on vesicular release during development. Neuron 85:145–158.

Neher E (2015) Merits and Limitations of Vesicle Pool Models in View of Heterogeneous Populations of Synaptic Vesicles. Neuron 87:1131–1142.

Nusser Z (2018) Creating diverse synapses from the same molecules. Curr Opin Neurobiol 51:8–15.

Ohtsuka T, Takao-Rikitsu E, Inoue E, Inoue M, Takeuchi M, Matsubara K, Deguchi-Tawarada M, Satoh K, Morimoto K, Nakanishi H, Takai Y (2002) Cast: a novel protein of the cytomatrix at the active zone of synapses that forms a ternary complex with RIM1 and munc13-1. The Journal of cell biology 158:577–590.

Renden R, Taschenberger H, Puente N, Rusakov DA, Duvoisin R, Wang LY, Lehre KP, von Gersdorff H (2005) Glutamate transporter studies reveal the pruning of metabotropic glutamate receptors and absence of AMPA receptor desensitization at mature calyx of held synapses. J Neurosci 25:8482–8497.

Schindelin J, Arganda-Carreras I, Frise E, Kaynig V, Longair M, Pietzsch T, Preibisch S, Rueden C, Saalfeld S, Schmid B, Tinevez JY, White DJ, Hartenstein V, Eliceiri K, Tomancak P, Cardona A (2012) Fiji: an open-source platform for biological-image analysis. Nature methods 9:676–682.

Schneggenburger R, Meyer AC, Neher E (1999) Released fraction and total size of a pool of immediately available transmitter quanta at a calyx synapse. Neuron 23:399–409.

Stanley EF (2016) The Nanophysiology of Fast Transmitter Release. Trends in neurosciences 39:183–197.

Taschenberger H, Leao RM, Rowland KC, Spirou GA, von Gersdorff H (2002) Optimizing synaptic architecture and efficiency for high-frequency transmission. Neuron 36:1127–1143.

tom Dieck S, Altrock WD, Kessels MM, Qualmann B, Regus H, Brauner D, Fejtova A, Bracko O, Gundelfinger ED, Brandstatter JH (2005) Molecular dissection of the photoreceptor ribbon synapse: physical interaction of Bassoon and RIBEYE is essential for the assembly of the ribbon complex. The Journal of cell biology 168:825–836.

tom Dieck S, Specht D, Strenzke N, Hida Y, Krishnamoorthy V, Schmidt KF, Inoue E, Ishizaki H, Tanaka-Okamoto M, Miyoshi J, Hagiwara A, Brandstatter JH, Lowel S, Gollisch T, Ohtsuka T, Moser T (2012) Deletion of the presynaptic scaffold CAST reduces active zone size in rod photoreceptors and impairs visual processing. J Neurosci 32:12192–12203.

Torres VI, Inestrosa NC (2018) Vertebrate Presynaptic Active Zone Assembly: a Role Accomplished by Diverse Molecular and Cellular Mechanisms. Mol Neurobiol 55:4513–4528.

Traynelis SF (1998) Software-based correction of single compartment series resistance errors. J Neurosci Methods 86:25–34.

Trussell L (1998) Control of time course of glutamatergic synaptic currents. Prog Brain Res 116:59–69.

Wagh DA, Rasse TM, Asan E, Hofbauer A, Schwenkert I, Durrbeck H, Buchner S, Dabauvalle MC, Schmidt M, Qin G, Wichmann C, Kittel R, Sigrist SJ, Buchner E (2006) Bruchpilot, a protein with homology to ELKS/CAST, is required for structural integrity and function of synaptic active zones in Drosophila. Neuron 49:833–844.

Wang T, de Kok L, Willemsen R, Elgersma Y, Borst JG (2015) In vivo synaptic transmission and morphology in mouse models of Tuberous sclerosis, Fragile X syndrome, Neurofibromatosis type 1, and Costello syndrome. Frontiers in cellular neuroscience 9:234.

Wang X, Hu B, Zieba A, Neumann NG, Kasper-Sonnenberg M, Honsbein A, Hultqvist G, Conze T, Witt W, Limbach C, Geitmann M, Danielson H, Kolarow R, Niemann G, Lessmann V, Kilimann MW (2009) A protein interaction node at the neurotransmitter release site: domains of Aczonin/Piccolo, Bassoon, CAST, and rim converge on the N-terminal domain of Munc13-1. The Journal of neuroscience : the official journal of the Society for Neuroscience 29:12584–12596.

Wang Y, Liu X, Biederer T, Sudhof TC (2002) A family of RIM-binding proteins regulated by alternative splicing: Implications for the genesis of synaptic active zones. Proc Natl Acad Sci U S A 99:14464–14469.

Wichmann C, Moser T (2015) Relating structure and function of inner hair cell ribbon synapses. Cell and tissue research 361:95–114.

Willsey AJ et al. (2018) The Psychiatric Cell Map Initiative: A Convergent Systems Biological Approach to Illuminating Key Molecular Pathways in Neuropsychiatric Disorders. Cell 174:505–520.

Xiao L, Michalski N, Kronander E, Gjoni E, Genoud C, Knott G, Schneggenburger R (2013) BMP signaling specifies the development of a large and fast CNS synapse. Nat Neurosci 16:856–864.

Zamponi GW (2016) Targeting voltage-gated calcium channels in neurological and psychiatric diseases. Nature reviews Drug discovery 15:19–34.

Zhang B, Seigneur E, Wei P, Gokce O, Morgan J, Sudhof TC (2017) Developmental plasticity shapes synaptic phenotypes of autism-associated neuroligin-3 mutations in the calyx of Held. Mol Psychiatry 22:1483–1491.

Zhen M, Jin Y (1999) The liprin protein SYD-2 regulates the differentiation of presynaptic termini in C. elegans. Nature 401:371–375.

